# Uncovering hundreds of exogenous and endogenous RNA viral RdRp sequences amongst uncharacterised sequences in public protein databases

**DOI:** 10.1101/2024.09.25.614983

**Authors:** K. Brown, A. E. Firth

## Abstract

Public databases of protein sequences, such as the National Center for Biotechnology Information (NCBI) Protein repository and UniProt, contain millions of proteins identified in samples from specific species but named as uncharacterised, hypothetical or unclassified due to a lack of information about their function. It has been demonstrated previously that many such sequences show high similarity to genes from RNA viruses, either due to viral infection of the original sample, contamination or endogenous viral elements (EVEs) integrated into the genome of the sample species. Many proteins from RNA virus discovery research are also deposited into these repositories but, for various reasons, can only be labelled as uncharacterised and classified taxonomically at a superkingdom or realm level. Sequences from protein repositories not labelled specifically as being derived from the RNA viral RNA dependent RNA polymerase (RdRp) protein are often used as negative controls when looking to identify viral RdRp sequences, so the presence of unlabelled viruses amongst these datasets is problematic.

In this study, we screened uncharacterised proteins from two large public protein repositories - NCBI Protein and UniProt - to identify sequences likely to be derived from RNA viral RdRp. 3,560 such sequences were identified, many derived from EVEs. Many previously unknown EVEs were identified and led to characterisation of additional, related sequences. For example, a group of orbivirus-like viruses infecting nematodes was uncovered which appears to have both ancient endogenous and circulating exogenous members. Many recent integrations of mito-like viruses into plant genomes were identified, indicative of current or recent RNA viral activity. In several taxonomic groups, the first example of an EVE, and in some cases the first example of any RNA virus, was uncovered. The large number of EVEs uncovered by this relatively small-scale search suggests that only a fraction of the true diversity of EVEs is currently known.

We also explore uncharacterised proteins further by providing provisional taxonomic annotations for RdRps which are currently only listed as members of the Riboviria realm. A number of sequences are identified which are indistinguishable from known, pathogenic viruses but are labelled as bacteria, seemingly as a result of mislabelling or contamination. Sequences which are not RNA viral but show some similarity to RdRp are also analysed, as a potential source of false positives in virus discovery research. Finally, recommendations are made for generating useful negative controls.

## Introduction

Over recent years, there has been an explosion in large scale screening of high throughput RNA-sequencing (RNA-seq) datasets for evidence of RNA viruses (for example, [1–16]. Most studies focus on identification via similarity to the RNA viral RNA-dependent RNA polymerase (RdRp) gene, the only gene which is conserved across all RNA viruses. Several tools have been published to aid in RNA virus detection in assembled RNA-seq data, for example RdRp-scan [17], PalmScan [18], VirSorter and VirSorter2 [19,20], and LucaProt [15].

Virus discovery research requires robust negative control sets of diverse, biologically derived sequences, known not to be derived from viral RdRp, to eliminate or reduce false positives. Approaches based on machine learning also require a training set of known non-viral sequences. An intuitive option is to use one of the many publicly available protein databases as a source of such sequences, with sequences annotated as non-RNA viral in origin selected as negative controls. Putative viruses that show higher similarity to any sequence in this negative control set than to any RNA virus are discarded as false positives.

Many successful virus discovery projects have used this approach, using similarity against proteins not annotated as RNA viruses from the NCBI non-redundant protein (nr) database [21] or the Pfam database [22] to eliminate potential false positives (for example, [1,2,4,8,10,14,17,23]. However, often these databases contain nucleotide or protein sequences which are derived from viral RdRp but are unannotated or mis-annotated. Also, many cellular genomes contain integrated endogenous viral elements (EVEs), derived from RNA viruses entering the genome of their host via chromosomal integration [24]. These are genuine host proteins but, given that they originate from RdRp, they do not represent false positives for RdRp detection software and therefore eliminating sequences similar to these proteins can be overly stringent.

Often, these unlabelled viral RdRp sequences will not be named as specific non-RNA viral proteins but instead have labels such as uncharacterised, unknown, putative or hypothetical protein or “domain of unknown function”. These labels are used when sequences are determined, either by the researcher or automatically, not to meet the criteria (based on homology, prediction of function or experimental evidence) to be assigned to a specific protein family. The sequences are often taxonomically assigned based on the organism which was the main source of the biological sample. However, biological samples ostensibly from a single organism will often also contain viral RNA from active infection or transcribed EVEs, in either the host, its environment or another source [13].

These unlabelled proteins, when derived from RNA viruses, hold great interest not only as as source of noise in virus discovery research, but also simply as a source of information about the RNA viruses present in biological samples. Despite virus discovery projects such as those listed above, unknown RNA viruses remain abundant and provide valuable insight into virus host range, transmission patterns and evolution. There are still species, and in some cases whole taxa, about the virome of which we know little. Taxa with poorly understood viromes tend to correspond to those with poorly annotated proteins, as under-studied species will have less data available to characterise genes. Therefore, uncharacterised proteins are likely to be a rich source of information about novel viruses infecting these species.

EVEs in particular are underexplored in comparison to exogenous viruses and are often dismissed as confounding factors in virus discovery research, but are an essential key to our understanding both the evolutionary history and the current distribution of RNA viruses. For example, very recent EVE integrations, such as the mitoviruses we have identified below in many plant genomes, are likely indicative of currently or recently circulating exogenous viruses. More directly, our identification of degraded reovirus-like EVEs in two nematode species led to our discovery of a related, likely active, virus in a distantly related nematode.

In this study, we present the results of large-scale screening of uncharacterised proteins from the National Centre for Biotechnology Information (NCBI) Protein [21] and UniRef [25] databases for RNA viral RdRp proteins, in order to clarify the origin of these proteins and facilitate virus discovery research. We characterise hundreds of new RdRp-like proteins, predominantly derived from EVEs. These results significantly enhance our understanding of the distribution of EVEs amongst under-studied species. We also provide taxonomic information for hundreds of known but unlabelled RNA viral RdRps, identify and list a number of likely mislabelled or contaminant-derived sequences, and characterise sequences which share near-significant similarity with RdRp. These findings are likely to serve as a valuable resource for future research on RNA viruses.

## Results

Over 300 million sequences labelled as unknown, unclassified or similar were identified for screening by searching the NCBI protein and UniRef100 databases - 75,483,361 from UniRef and 233,325,610 from NCBI.

Profile HMMs (pHMMs) have been previously shown to be a sensitive and accurate technique for detecting viral proteins [16,26], therefore a pHMM-based approach was used to initially detect candidate RdRps. Pre-existing profiles from two recent publications [16,26] and from the Pfam [22] database were used to ensure a comprehensive search.

An initial 92,741 uncharacterised sequences had significant matches (HMMER score ≥ 25) against these pHMMs. After additional filtering (described below), 3,560 sequences were identified as likely to be RNA viral RdRp - 1,609 classified as Viruses/Riboviria (but not Orthornavirae) in their source database. 1,460 as Eukaryota, 251 as Bacteria, 210 as synthetic, unidentified or vector and 30 as metagenomic (Figure 1).

**Figure 1:**
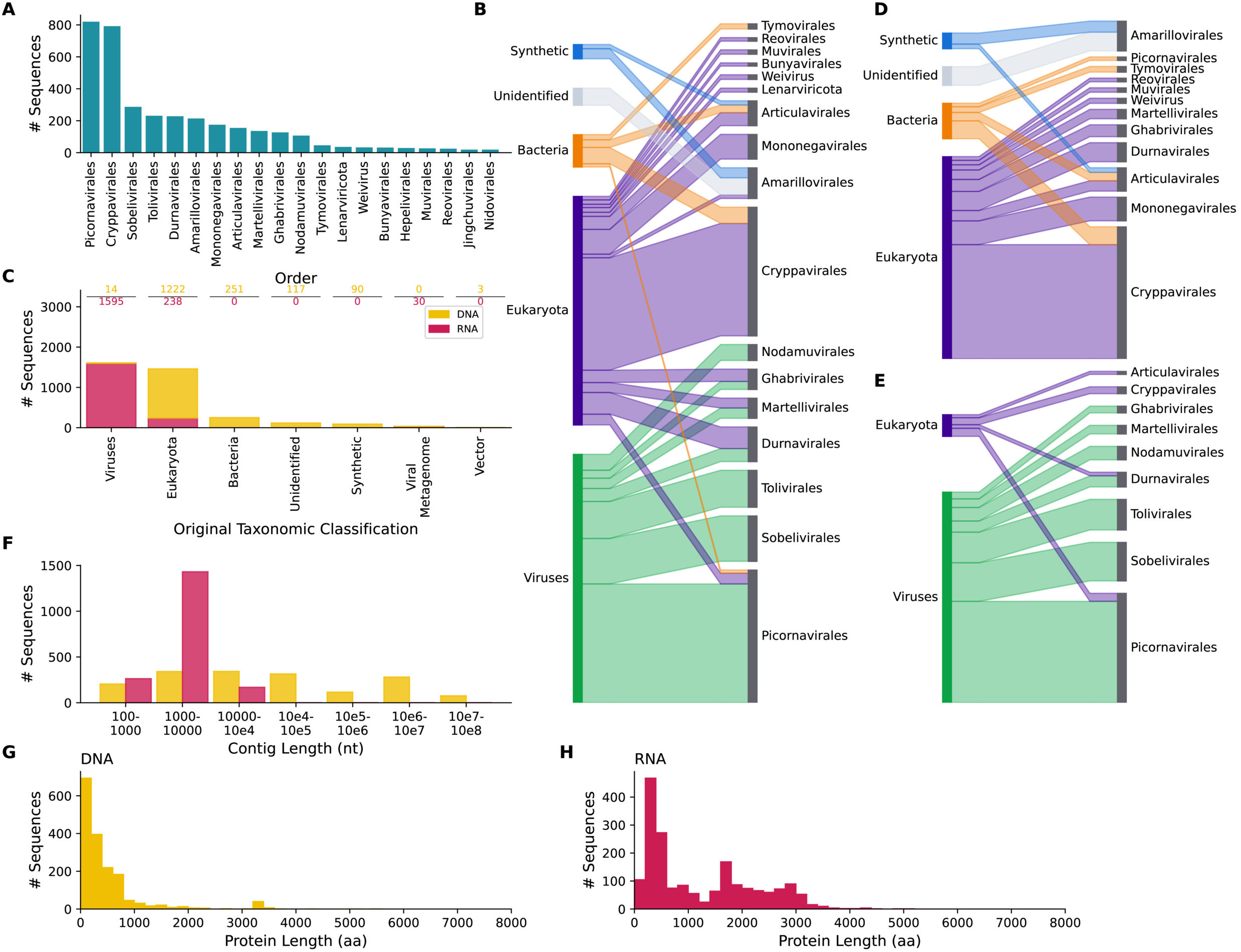
Summary of Initial Results. (A) Bar chart showing the number of RdRp-like uncharacterised proteins classified via phylogenetic analysis into each of the ICTV orders into which at least 10 proteins were classified. (B) Sankey plot comparing the classification of all identified RdRp-like uncharacterised proteins based on their NCBI taxonomy ID versusthe viral order into which they were assigned. (C) Bar chart comparing the number of uncharacterised sequences with each classification based on their NCBI taxonomy ID for those labelled as having a DNA (yellow, top) or RNA (pink, bottom) source, numbers above bars correspond to bar height. (D, E) Sankey plots as in (B) but for DNA source sequences (D) or RNA source sequences (E) only. (F) Histogram indicating the length distribution of source contigs categorised by their original taxonomic classification. (G, H) Histogram indicating the protein sequence length distribution of uncharacterised proteins categorised by their original taxonomic classification.

### Filtering Reverse Transcriptase

Of the 92,741 sequences identified with the original set of RdRp pHMMs, a large majority (90%) matched one of two profiles; the Pfam profiles PF05919 (Mitovirus RNA-dependent RNA polymerase) and PF00680 (Viral RNA-dependent RNA polymerase). However, inspection of these sequences revealed that many have higher homology to the retroviral reverse transcriptase (RT) protein (and, correspondingly, retrotransposon RT) than to RNA viral RdRp. RT sequences are appropriate negative controls for RNA virus discovery experiments, as RdRp and RT share distant homology [27] and RT is a common source of false positives.

In order to identify RT-like hits amongst the virus-like unclassified proteins, a phylogenetic analysis was performed on the sequences included in input alignments used to generate the two Pfam profiles (Figure S1). In both cases, a monophyletic group, primarily of named RdRp proteins, formed, clearly distinct from RT sequences. A few scattered sequences tagged as other proteins also fell into the RdRp clades. For PF00680, A0A1B0C5W8, labelled as the phospholipid scramblase protein (PLSCR2) of the tsetse fly *Glossina palpalis,* was in the RdRp-like cluster (Figure S1A). For PF05919, A0A371IGY1, labelled as the velvet bean *Mucuna pruriens* NADH-ubiquinone oxidoreductase chain 6 (ND6) and A0A5A7QJR6, labelled as red witchweed *Striga asiatica* Photosystem I P700 chlorophyll a apoprotein A1 (PSAA), were in this group (Figure S1B). In all three cases, BLAST searches showed different sections of the sequences were highly similar to either RdRp or to eukaryotic proteins, suggesting either endogenisation of viral sequences or misassembly. A number of unclassified sequences fell into the RdRp-like group in both cases (Figure S1).

New pHMMs were generated by dividing the Pfam profile input sequences into RdRp-like and RT-like clusters. All hits against PF05919 and PF00680 were compared to these clusters individually, and designated as RdRp or RT depending on the highest score. Removing sequences with higher similarity to RT than RdRp left 9,249 putative viral sequences.

### Identifying RdRp

Several additional analyses were performed to determine if sequences were truly RNA viral in origin. Sequences were assigned using phylogenetic analysis to one of 330 viral operational taxonomic units (OTUs). The OTU assigned to each sequence is listed in Supplementary Table 1 and individual trees for each OTU are available in the Supplementary Data. Sequences were screened using DIAMOND BLASTP [28] to ensure that every sequence was either more similar to a named RNA viral protein than to any other named protein, matched only other uncharacterised proteins or matched nothing.

The predicted protein structures of all sequences were compared to PDB virus profiles using FoldSeek [29], PalmScan [18] was used to identify core RdRp regions and BLASTP [30,31] was used to make an additional sequence similarity comparison to known RdRp sequences.

Sequences were required to score highly on at least two of the following criteria to be classified as RdRp-like: 1) A FoldSeek bit score of >100 against an RdRp profile, 2) an identified RdRp core using PalmScan, 3) a BLAST bit score of >50 against a known RdRp sequence, 4) a branch length of less than 1.5 separating the sequence from the nearest known RNA virus in phylogenetic analysis and 5) a HMMER score >50 against an RdRp profile. After this filtering step, 3,560 sequences remained. Results from running these analyses are listed in Supplementary Table 1.

An important caveat is that, as the NCBI protein database is not curated, many of the sequences identified here will not be unique, but will be the result of either multiple projects sequencing the same genome, chromosome or other sequence or overlapping contigs or scaffolds from the same study. Therefore, “sequences” in this context refers to individual NCBI protein / UniProt records, rather than unique biological entities.

### RdRp Diversity

The largest proportion of sequences identified here, 820 sequences, are Picornavirales, a large order of positive sense RNA viruses, followed by the Cryppavirales (792 sequences), another positive sense order containing only one family, the Mitoviridae [32] (Figure 1A, 1B). The distribution of sequences between orders was very distinct depending on the assigned taxonomic origin of the original database entry (Figure 1B). The majority of the Picornavirales, Sobelivirales, Tolivirales and Nodamuvirales sequences were already classified as viral (but not RNA viral) in their source databases, while most Cryppavirales, Mononegavirales, Articulavirales and Bunyavirales were classified as eukaryotic.

As RNA viruses only use RNA as their genetic material, it was expected that RdRp-like sequences would be largely derived from RNA, for example sequences assembled from RNA-sequencing datasets. However, surprisingly, 48% of the RdRp-like proteins are labelled in their source database as originating from genomic DNA (Figure 1C). Although all proteins are translated from RNA, many protein database records are inferred open reading frames (ORFs) identified automatically via gene prediction on DNA sequences, using tools such as Augustus [33]. Therefore, the protein records link directly back to the DNA records without mRNA intermediates.

There are three possible sources of RNA viral material in DNA sequences – contamination, mislabelling and endogenisation of RNA viruses in host genomes. In the case of contamination or mislabelling, we would expect the DNA-source RdRps to resemble those with an RNA source. However, EVEs differ from circulating exogenous viruses in a number of ways.

Firstly, the taxonomic distribution of EVEs differs from that of exogenous viruses. The eukaryotic DNA putative RdRps (EDRdRps) examined here most often originate from plants (656/1,222 sequences) and arthropods (274/1,222 sequences). Arthropods are known to have EVEs particularly from the Mononegavirales, Martellivirales and Durnavirales [24] while plants have many Cryppavirales and Durnavirales [13,34,35]. Arthropod and plant EVEs are seemingly rarely derived from the Picornavirales, an abundant exogenous order amongst these hosts [36]. The EDRdRps examined here are most commonly classified as Cryppavirales (52% of sequences), Mononegavirales (11%) and Durnavirales (9%) (Figure 1D). Meanwhile, Picornavirales are rare, with only eight eukaryotic sequences identified. These percentages differ from the RNA-source RdRps, amongst which Picornavirales make up 42% of sequences, while Cryppavirales, Mononegavirales and Durnavirales are 3%, 2% and 6% respectively (Figure 1E). Therefore, the taxonomic distribution of the EDRdRps is more consistent with that of EVEs than of RNA viruses.

Secondly, EVEs will be integrated into host chromosomes, so the source nucleic acid sequences, either chromosomes, scaffolds or contigs, would be expected to often be of a chromosome-like length, i.e. on a megabase scale. Meanwhile, the longest known RNA viral genome is a 41.1kb Nidovirus [37], so RNA viral transcripts are substantially smaller. Here, the DNA-source RdRps have a mean source sequence length of 14.7Mb, compared to 5,402nt for the RNA-source RdRps sequences, an increase of more than 2,500 fold (Figure 1F). The majority of RNA source sequences are between 1kb and 10kb in length. This again suggests that the eukaryotic DNA-source sequences generally represent EVEs.

Thirdly, once EVEs enter the host genome they are often no longer subject to selection pressure to maintain an intact open reading frame [38], so the ORFs will be short and degraded compared to those of exogenous viruses. Therefore, the protein sequences of EVEs should be shorter, on average, than those of RNA viruses. Accordingly, the DNA-source RdRps here have a mean length of 482 amino acids, while the RNA-source RdRps have a mean length of 1,228 amino acids (Figure 1G, 1H).

Taken together, these observations suggest that EVEs are numerous amongst our DNA-source RdRp sequences.

As prokaryotes have no known EVEs, the bacterial RdRp-like elements from DNA sequences are likely to represent mislabelling or contamination, these will be discussed separately below.

### EVEs derived from Positive Strand RdRps

#### Cryppavirales

A high proportion (52%) of the identified EDRdRps are members of the Cryppavirales order, which contains one family, the Mitoviridae (Figure 1D) [32]. Almost all of these sequences (637/640) are from plants from the Streptophyta clade; the remaining three are fungal. With the exception of one sequence in white spruce (*Picea glauca*), all plant sequences are from flowering plants of the Magnoliopsida class.

The plant DNA-source Mitoviridae have a mean ORF length of 305 amino acids, compared to 774 amino acids for ICTV classified Mitoviridae (Figure 2A). This suggests that these sequences may be EVEs. Many plant genomes are known to contain mitovirus derived insertions, particularly in the mitochondrial genome [39–42]. Mitovirus-like EVEs in plants are thought to have originally integrated into plant mitochondrial DNA, but then, in some cases, been transferred into the nuclear genome [35]. Accordingly, approximately one quarter (170/637) of our identified plant DNA-source mitoviruses are from sequences labelled as mitochondrial.

**Figure 2:**
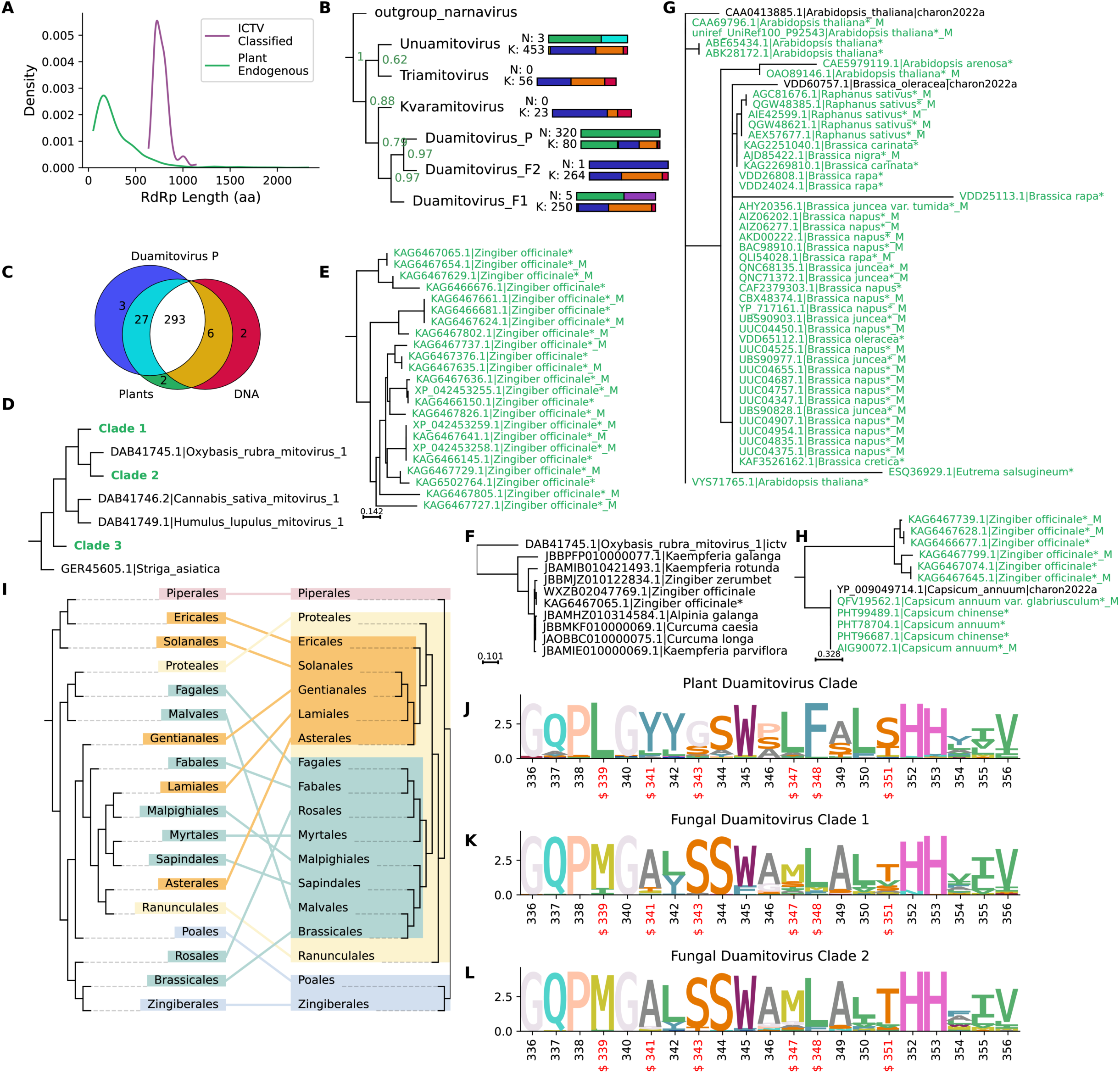
Mitovirus-like EVEs. (A) Density plot showing the length distribution of plant DNA source uncharacterised proteins classed as Mitoviridae in comparison to proteins from ICTV classified Mitoviridae. (B) Summarised phylogenetic tree showing the distribution of the Mitoviridae genera (full phylogeny is available as Supplementary Figure 4). The tree has been simplified to show only these genera and exclude sequences which didn’t fall clearly into the clade for any genus. Bars at node tips show the number of sequences in each group, the top bar (N, new) is the number of newly identified sequences and the bottom bar (K, known) the number of previously annotated sequences. Bar colours show the taxonomic origin of the sequences as follows: green, plants; orange, viruses; pink, metazoa; blue, metagenomic; purple, fungi; cyan, SAR. (C) Venn diagram showing the overlap in sequences assigned to the Duamitovirus P clade, sequences identified in plants and sequences with a DNA source. (D) Simplified subsection of the Mitoviridae phylogenetic tree showing the relationship between the clades expanded in the subsequent subfigures. (E) Expansion of clade 1 from the phylogeny section shown in (D). Sequences marked with asterisks and with green text are newly identified, sequences with the suffix _M are from the mitochondrial genome. (F) Phylogeny incorporating additional sequences from members of the Zingiberoideae family identified by further analysis of the sequences in clade 1. (G) Expansion of clade 2 from the phylogeny section shown in (D). Labels are as for (E). (H) Expansion of clade 3 from the phylogeny section shown in (D). Labels are as for (E). (I) Tanglegram comparing the phylogenetic relationships of Magnoliopsida plants, generated manually based on [45] and those of the most common clade of mito-like viruses identified in each taxon (approximate maximum likelihood generated with FastTree2 [84]). Colours correspond to plant taxa. (J-L) Sequence logos for RdRp motif B from the Duamitovirus P clade (J) and fungal Duamitovirus clades F1 (K) and F2 (L). Logos for RdRp motifs A to C are available in Supplementary Figure 5. Variable positions are labelled in red and with “$”. Positions are relative to the full length alignment available in the Supplementary Data.

An additional 48 plant and 4 non-plant RNA-source Mitoviridae RdRps were also identified. 36 of these RNA-source sequences are mRNA sequences of mitochondrial proteins (eukaryotic mRNA putative RdRps, EMRdRps), so they most likely represent transcribed sequences from the integrated RdRps, rather than transcripts from exogenous viruses.

Given that many EVEs are degraded and partial sequences are common, to ensure the accuracy of the phylogeny and avoid sequences clustering by which region of RdRp is present, only sequences fully overlapping the same 30 amino acid region and with a length of at least 100 amino acids after trimming were selected for phylogenetic analysis. This left 328 plant sequences (Figure S2). These sequences represent most plant orders in the original dataset, however EVEs from some, most notably the Asparagales, were too degraded to include (Figure S2). An additional 5 sequences from non-plant hosts overlapping this region were also included (Figure S3).

The basic structure of the phylogeny generated using these sequences, along with known mitovirus sequences from ICTV, RefSeq and previous virus discovery projects is shown in Figure 2B. The full phylogenetic tree is shown in Figure S4. The majority of the sequences appear to be newly identified, endogenous members of the Mitoviridae family.

The major taxonomic groups of mitoviruses were apparent in our analysis, with the ICTV-classified Duamitovirus, Unuamitovirus and Triatomitovirus genera forming monophyletic clades together with many unclassified viruses and virus sequences from other virus discovery projects (Figure 2B, Figure S4). The Kvavomitovirus genus only contains one classified sequence, however it clustered separately from the other genera and together with a small number of metazoan and metagenomic sequences from a previous study which classified many sequences from publicly available RNA-seq datasets using the Serratus architecture [42]. The phylogenetic relationship between genera is not well-established, however here the Triatomitovirus and Unuamitovirus genera are sister clades, with the Duamitovirus genus more distantly related and sharing a more recent common ancestor with the Kvaromitovirus group. A number of previously annotated sequences did not fall into any of the established genera, one of our newly identified sequences also did not, from an environmental sample from *Dinobryon* algae (CAF8515450.1) (Figure S4). This sequence appears to represent a new example of Telchines mito-like virus, identified previously in algae [43].

Almost all of the RdRps (320/333) in our study fell into a single, large clade within the Duamitovirus genus, designated here as the Duamitovirus Plant (P) clade (Figure 2C). All but three of these sequences were from plants and 293 had a DNA source. This clade also included 80 sequences from other studies (Figure 2B), including all of the eight ICTV classified Magnoliospida duamitovirus sequences [40], plus one classified duamitovirus, from an aquatic fern, *Azolla filiculoides*. The Duamitovirus P clade is distinct from two other large clusters within the Duamitovirus genus, which contain only eight of our identified RdRps. These clades contain many named (ICTV-classified and unclassified) viruses of plant pathogenic and symbiotic fungi, plus numerous sequences from virus screening projects against metagenomic datasets.

Within the P clade, many clusters of RdRp-like sequences are apparent. A small example subsection of the phylogeny is summarised in Figure 2D, many other similar regions are present within the P clade. Within this subsection, clade 1 (Figure 2E) contains newly identified sequences, all of which are from the ginger plant (*Zingiber officinale*). This clade likely represents a previously unknown RdRp integrated into the genome of *Z. officinale*. The sequences from this clade are all from the same ginger genome sequencing project [44] (assembly Zo_v1.1). Therefore, for verification, a BLAST search against the online server was used to identify other contigs from members of the ginger family, Zingiberaceae, which contain this EVE. Highly similar ORFs (78-99% identity) were identified in a contig from the other *Z. officinale* reference genome available on GenBank (assembly ASM1131758v2) and in seven other species in this family (Figure 2F), including members of both the Zingiberoideae and Alpinioideae, the only subfamilies with contigs currently available through GenBank. This clade therefore represents a novel EVE which is likely to be widespread in different members of this plant family.

Clade 2 contains novel EVEs from a number of Rosid plants, members of the genera *Brassica*, *Raphanus*, *Arabadopsis* and *Eutrema* (Figure 2G). Two sequences from the RdRp-Scan database are also in this clade. Particularly between members of the *Brassica* and *Raphanus* genera, there is almost no variation between these sequences. This suggests that these EVEs are either very recent insertions, and therefore have had little time to accumulate mutations in their hosts, and therefore that related exogenous viruses are currently circulating or that there is some kind of selection pressure maintaining the ORF.

Similarly, clade 3 (Figure 1H) contains six 100% identical sequences, from two species of chilli pepper - *Capiscum chinense* and *Capiscum annuum*. These sequences are from the mitochondrial genome (QFV19562.1, AIG90072.1) and chromosome 6 (PHT78704.1) of *C. annuum* and chromosome 12 (PHT99489.1) and an unplaced scaffold (PHT96687.1) of *C. chinense* (plus one additional sequence from the RdRp-scan database). The perfect conservation of these sequences again suggests either very recent integration events or selection pressure.

The tanglegram in Figure 2I shows the relationship, at taxonomic order level, between the host phylogeny [45] and the biggest group of endogenous RdRps identified in each order. It is not possible to recapitulate the host phylogeny using the mitovirus sequences. This implies that the viruses became endogenous comparatively recent and have not been present throughout the evolution of their hosts. There are some similarities between the trees, for example the Malpighiales, Myrtales and Sapindales, all of which are Rosids, cluster together, as do the Ericales and Solanales, which are both asterids. However, overall the trees are not highly similar, therefore, it is likely that the insertion of these elements into the genomes of their hosts occurred after the divergence of the host orders from a common ancestor. Where closely related hosts do share closely related insertions, such as those in the Ericales and Solanales, or Malpighiales and Sapindales, this is likely to be because more similar hosts are more likely to be susceptible to the same viral strains, rather than because the insertions occurred in a common ancestor.

The sequence logos in Figures 2J-L and Figure S5 show the conservation of key residues in different subsets of mitoviruses. 13 residues in motifs A to C are conserved (defined here as a bit score ≥ 3) in ICTV classified mitoviruses and in all six major clades of the phylogenetic tree. These include the key residues essential to RdRp function [46], the N-terminal aspartates Dx_4–5_D are apparent at positions 286 to 292 in motif A as is the GDD element at positions 374 to 376 in motif C (Figure S5). The level of similarity between the Duamitovirus P clade, which is largely endogenous, and the other clades, suggests again that the endogenous mitovirus-like elements are relatively recent insertions and have not fully degraded.

However, the P clade does clearly differ from the other Duamitoviruses at some positions, which are concentrated in motif B (Figure 2K-L). The conservation patterns of the P clade are consistent with those apparent in plant mitovirus EVEs examined previously [35]. These changes seem to be largely specific to Duamitoviruses infecting plants as opposed to plant pathogenic fungi and to have been acquired in the lineage leading to the Duamitovirus P clade. Most of the changes are between amino acids with similar properties (methionine to leucine at positions 339 and 347, serine to threonine at position 351), however the glycine to serine change at position 343, phenylalanine to leucine change at 348 and particularly the tyrosine to alanine change at position 341 may be more impactful. The methionine at position 339 is somewhat characteristic of the Lenarviricota, present in at least the Botourmiaviridae, Fiersviridae and Narnaviridae, while most ssRNA+ viruses have a strongly conserved serine at this position [47]. However, of the 299 Magnolopsida sequences which have coverage at this position, 290 have a leucine residue. These same families of Lenarviricota lack the highly conserved Asparagine residue at position 348 seen in other ssRNA+ viruses, which has been demonstrated to be essential for NTP binding [46], and instead have leucine (Narnaviridae, Botourmiaviridae) or glutamic acid (Fiersviridae) at this position [46,47], while in the P clade phenylalanine is most common.

#### Martellivirales

The other highly represented group of eukaryotic EVEs derived from positive strand RNA viruses is the Martellivirales order, with 60 sequences (53 EDRdRps and 7 EMRdRps). These sequences have a mean length of 549 amino acids, compared to 1,674 amino acids for ICTV classified Martellivirales. The majority of the sequences are fungal (40/60), while 16 are from arthropods. Most (37/40) of the fungal sequences are near-identical Virgavirus-like insertions from the pathogenic fungus *Rhizopus arrhizus* (Supplementary Data: Tree Martellivirales_14), these seem to be endogenous insertions related to the exogenous virgavirus of *Rhizopus* identified previously [48].

The arthropod sequences are more scattered, with several single sequences which resemble members of the Virgaviridae and Kitaviridae. Such plant viruses are known to occur as EVEs in arthropod genomes [49]. The sequences found here fall within the diversity of known Kitaviridae and Virgaviridae species. Two small example clusters of arthropod EVEs are shown in Figure S6A. The first cluster contains five EMRdRps from the western flower thrip *Frankliniella occidentalis*, likely to represent a novel group of EVEs. The closest relative is Eriocheir sinensis kita-like virus, an recently identified exogenous Kitaviridae-like virus from another arthropod, the Chinese mitten crab [50]. The second cluster is found in two Lepidopteran species, the moth *Arctia plantaginis* and the butterfly *Pieris macdunnoughi,* closely related to an exogenous butterfly virus, Polyommatus icarus virga like virus 2 [8].

### EVEs derived from Negative Strand RdRps

#### Mononegavirales

Amongst the eukaryotic RdRps resembling negative-strand RNA viruses, members of the order Mononegavirales were the most abundant, with 161 sequences. 142 of these are EDRdRps and 11 EMRdRps, the remaining eight are from assembled RNA-seq datasets, and therefore cannot be assumed to be endogenous. The Mononegavirales sequences are primarily from arthropods (96 sequences) and from a specific member of the Acanthocephala group of parasitic worms, *Pomphorhynchus laevis* (18 sequences).

Figure S6B shows the phylogeny of the 16 of the 18 *P. laevis* sequences. A sequence from the Serratus screening project [11], also from *P. laevis,* is also within this group, as this project used RNA-sequencing data these RdRps must at least sometimes be transcribed. No viruses infecting this host have otherwise been previously described. These sequences cluster close to the Cytorhabdovirus genus of Rhabdoviridae. This genus is generally considered to consist of plant viruses, however, several viruses isolated from arthropods are ICTV-classified members of this group, including viruses from Crustacea, a group used as intermediate hosts by *P. laevis* [51]. All of the *P. laevis* RdRp-like sequences are from unplaced contigs from a single genome project [51], the only assembly available for any member of the Acanthocephala, so it is not clear how widespread this putative EVE is within this group. It is also not possible to say definitively that these contigs are from the specified host, rather than another species present in the sample, especially given that the assembly is derived from an intestinal sample. However, the sequences are relatively divergent (mean 60% identity), fragmented in different ways and contain multiple insertions and deletions, which suggests there are multiple insertions in the host genome and that at least some of the EVEs are ancient.

Amongst the 96 arthropod EVEs identified, 60 are from species of Lepidoptera and are widely distributed through the Mononegavirales phylogeny, with clusters related to the Nyamiviridae, Xinmoviridae, Rhabdoviridae and Artoviridae. Some of the Rhabdoviridae-like sequences are shown in Figure S6C. Spodoptera frugipdera rhabdovirus is a known, transcribed, EVE of the fall armyworm *Spodoptera frugipdera* [52]. Sequences identified in three contigs from the wood tiger moth *Arctia plantaginis* closely resemble the RdRp of this virus. A second cluster of more distantly related sequences was found in the diamondback moth, *Plutella xylostella,* resembling transcribed sequences identified in the almond moth, *Cadra cautella* [16].

A small cluster of RdRp-like EVEs were identified in *Perkinsus olseni*, a parasitic species of single-celled protists (Figure S6D). Nine sequences were found across three genome assemblies for this species. These are the first EVEs which have been identified in the Perkinsozoa phylum and no RNA viruses have been described to date infecting this phylum except a single fragment of a toti-like RdRp identified in *Perkinsus chesapeaki* [26]. The *P. olseni* sequences we identified here are divergent from any known virus family. The closest related group is the Lispiviridae (Figure S6D), however the highest scoring BLAST hit against any ICTV-classified virus shares only 23% identity (KAF4688156.1 against YP_010800584.1 from Copasivirus ivindoense, Isopteran arli-related virus OKIAV103). These RdRp fragments are likely to be derived from a unique, monophyletic group of Mononegavirales. The sequences are fragmented and from different parts of the RdRp gene, however they can be combined into a 1272 amino acid consensus, as shown in Figure S7 (alignment available as Supplementary Data). Across the aligned regions, excluding gaps, there is a minimum of 82% identity between sequences, and a mean of 91%. A HHpred [53] search confirms similarity along the majority of this consensus (976/1272 amino acids) to the “Mononegavirales RNA dependent RNA polymerase” (PF00946.23) and “Mononegavirales mRNA-capping region” (PF14318.10) Pfam records.

#### Articulavirales

67 eukaryotic DNA and mRNA-derived sequences were also identified in the *Articulavirales* order, all of which were closely related to the Orthomyxoviridae (Supplementary Data: Tree Articulavirales_2). RdRp-like sequences in this group were identified in genomic sequences from many arthropod groups including the Coleoptera, Lepidoptera, Heteroptera and Diptera. The subset shown in Figure S6E are from a number of Lepidopteran species. Orthomyxoviridae EVEs have previously been identified in arthropods [54,55] and it appears that Orthomyxoviridae commonly endogenise in this taxon.

### EVEs derived from Double Stranded RNA Viruses

#### Durnavirales

107 EDRdRps and 19 EMRdRps were identified in the Durnavirales order of dsRNA viruses. 71 of these sequences are from members of the Mucoromycota division of fungi. Of these 71, 64 fall within a single clade, shown in Figure 3A, and are found in the Mortierellaceae family of fungi. The mean length of sequences in this clade is 646 amino acids, compared to 1,495 amino acids for ICTV classified Durnavirales (Figure 3B). The closest ICTV classified virus to these putative EVEs is Curvularia thermal tolerance virus, a member of the Curvulaviridae family of viruses. Its host, *Curvularia,* is a member of the Pleosporaceae family of fungi.

**Figure 3:**
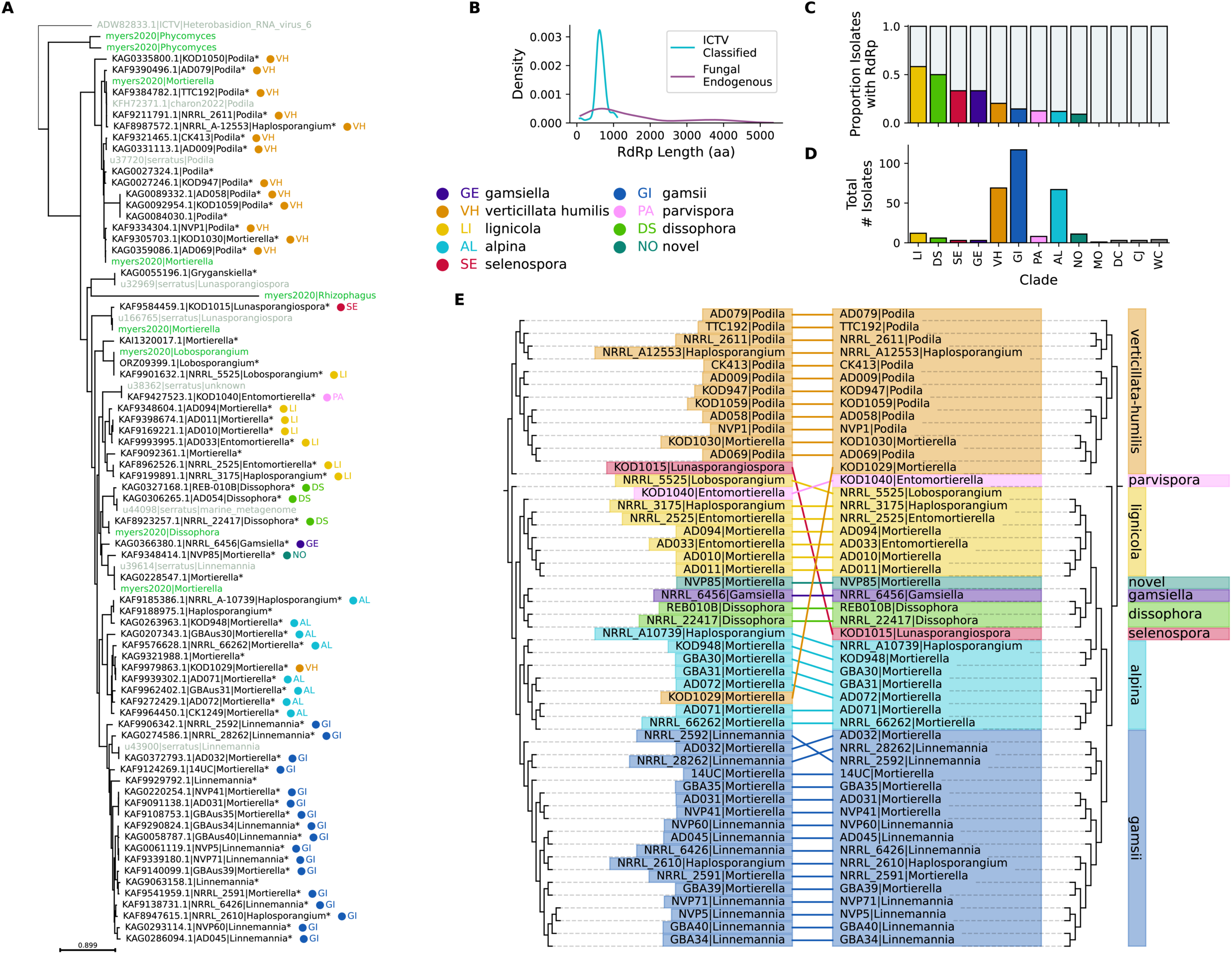
Durnavirales-like EVEs. (A) Approximate maximum likelihood [84] phylogenetic tree showing the newly-identified clade containing EVEs from the Mortierellaceae family of fungi and related sequences. The full phylogeny is available in the Supplementary Data. Node tips marked with a dot are from [57] and are labelled with the cluster into which the host species were classified by these authors (named hereafter as Vandepol clusters), as shown in the legend. These sequences are labelled with their GenBank accession, the ID originally assigned to the fungal isolate and their GenBank genus. Newly identified sequences are marked with an asterisk (*). Sequences with the prefix Myers2020 and shown in green are fungal viruses from the unclassified group described by [56]. Sequences shown in grey are from previous large scale virus discovery research projects - either Serratus [11] (labelled as serratus) or RdRp-Scan [26] (labelled as charon2022). (B) Density plot showing the length distribution of fungal DNA source uncharacterised proteins classified in this clade in comparison to proteins from ICTV classified Durnavirales. (C) The proportion of isolates from of each of the Vandepol clusters with contigs showing similarity to RdRp. (D) The total number of isolates sequenced in each of the Vandepol clusters. (I) Tanglegram comparing the phylogenetic relationships of Mortierellaceae plants, based on [57], and those of the most common clade of durna-like viruses identified in each taxon. Colours correspond to Vandepol clusters and prefixes to the isolates labelled in the same publication.

Our newly identified Mucoromycota group forms a separate monophyletic group to this virus and its relatives. Several previously identified but unassigned viruses [56] in Mucoromycota do fall within this newly identified group (Figure 3A). Several Serratus sequences from the Mortierellaceae and one from a marine metagenomic dataset are also within this group, as is one sequence identified by RdRp-Scan [11,26].

Of the sequences we identified in this clade, almost all (53/64) are from the same publication, which investigated the phylogeny of the Mortierellaceae [57]. No viruses or EVEs were discussed in this publication, which focused on host phylogenetics. However, of the 316 isolates sequenced in this paper, 53 included contigs with members of this putative EVE family. These authors divide their sequences into 14 clusters, which they propose as genera to replace the seven which were recognised at the time of their publication. The frequency of the EVE is highly variable between these clusters, over 50% of the “lignicola” group have a contig containing the insertion, while no insertions were present in contigs from the modicella, dichotoma, cystojenkinii and wolfii capitata groups (Figure 3C). These groups do have relatively low representation in the dataset, so it is possible that the EVEs would be present if more samples were included (Figure 3D). It may also be the case that the more represented host groups have more copies of the EVE, which will therefore have more coverage amongst the sequencing reads.

Figure 3E compares the host phylogeny from [57] with the relationships between the EVEs. The two trees are much more similar than those for the Mitoviruses (Figure 2I) and the majority of the host tree can be recapitulated using the EVE tree. This suggests either a much longer standing relationship between these viruses and their hosts, with EVEs inserted prior to the last common ancestor of the members of this family, or a very strong host specificity of the viral strains. The sequences within the clade are relatively diverse, with a mean similarity of 66%, this again suggests that these EVEs may be ancient.

47 of the 55 remaining putative eukaryotic Durnavirales EVEs are found in arthropods. These sequences form multiple small clusters, distributed amongst various families of Durnavirales. Two such clusters are shown in Figures S6F and S6G. Four sequences were identified in the seven-spot ladybird *Coccinella septempunctata* which cluster with the Partitiviridae (Figure S6F). On further examination, these sequences are predicted transcript isoforms from a single host gene on chromosome 2, zinc finger protein 64-like LOC123308761, so they represent a single EVE insertion event.. The structure of this locus consists of six transcript isoforms, four of which contain this insertion, with a long central intron. The four isoforms with the EVE encode a single ORF at the 5’ end, upstream of the intron (Figure S8, alignment available in Supplementary Data). The two isoforms without the EVE also encode a single ORF, at the 3’ end, downstream of the intron (Figure S8). The arrangement of these predicted isoforms suggests that this EVE is not part of the gene, but rather that the intron is mislabelled and only the 3’ ORF is valid. There is some homology between positions 1651 to 1791 and positions 13962 to 14071 of this locus, which could have led to incorrect splice junction prediction (Figure S8).

A second cluster of Durnavirales is seen in sequences from the South American giant ant *Dinoponera quadriceps*, again clustering with the Partitiviridae (Figure S6G). Similarly to the *C. septempunctata* sequences, these proteins are from predicted transcript isoforms from the same host gene, uncharacterised LOC106741364. This genome is not assembled to chromosome level, however this locus is on a 4Mb scaffold, so it is on a chromosome-like scale. All of the transcript isoforms predicted at this locus have the EVE as their one predicted ORF, which is in the middle of a longer transcript. Online BLASTX of this transcript shows no similarity to proteins other than the EVE, however there is some limited nucleotide similarity to other *D. quadriceps* transcripts on either side of the EVE. It is not possible to determine from the data if this is a real transcript interrupted by the EVE or an annotation error.

#### Reovirales

Reovirales EVEs are not particularly well-known, but have been previously identified in mosquitoes [58]. We found a number of EVEs related to the Reovirales, specifically to proposed (but not ICTV classified) members of the Orbivirus genus, in a well-supported clade containing primarily exogenous insect viruses (Figure S6H). Most of the EVEs we identified in this clade amongst our uncharacterised proteins are also from insects, with two sequences from the Edith’s checkerspot butterfly (*Euphydryas editha*), one from the Mexican paper wasp (*Mischocyttarus mexicanus*), one from the cabbage stem flea beetle (*Psylliodes chrysocephala*) and five from the Asian citrus psyllid (*Diaphorina citri*). These diverse insect insertions suggest that endogenous Reovirales are more widespread amongst insects than has previously been thought, especially within this specific clade.

Interestingly, five EVE sequences were also found within this clade amongst uncharacterised proteins from two nematode species - the parasitic roundworms *Brugia pahangi* and *Brugia malayi* (Figure S6H). These worms are from the Onchocercidae family of filarial nematodes. Both are pathogenic, *B. malayi* causes lymphatic filariasis in humans and *B. pahangi* causes filarial disease in cats and dogs, with occasional zoonotic transmission to humans [59]. Both are vectored by various species of mosquito [60].

On further interrogation based on our initial finding in *Brugia*, using TBLASTX against the reference genomes for these two species (NCBI accessions GCA_012070555.1 and GCA_000002995.5), several regions resembling reoviral RdRp were identified. In both species, it was possible to identify an approximately 2,400 nt region of chromosome X, 1,600 nt region of chromosome 4 and 700 nt section of chromosome 2 with recognisable homology to the orbivirus-like Reovirales (Figure 4A). There are no ICTV classified viruses within this clade, so the full RdRp sequence of Hubei lepidoptera virus 4 (KX884628.1) was used as a full length reference. This sequence is 4,041 nt in length, so the chromosome X region is approximately 60% full length. The sequences were found to share far higher similarity with regions from the same chromosome in the other *Brugia* species than with insertions on other chromosomes within the same species. The chromosome X sequences from *B. pahangi* and *B. malayi* are 96% identical to each other on a nucleotide level, the chromosome 4 sequences 97% identical and the chromosome 2 sequences 98% identical (Figure 4A, 4B). Meanwhile, the three *B. pahangi* sequences share 38-62% nt identity with each other and the three *B. malayi* sequences 57-79%. All of the identified regions are highly degraded and do not contain the expected long RdRp ORF, instead they contain many stop codons and frameshifts, typical of comparatively ancient endogenous insertions [38].

**Figure 4:**
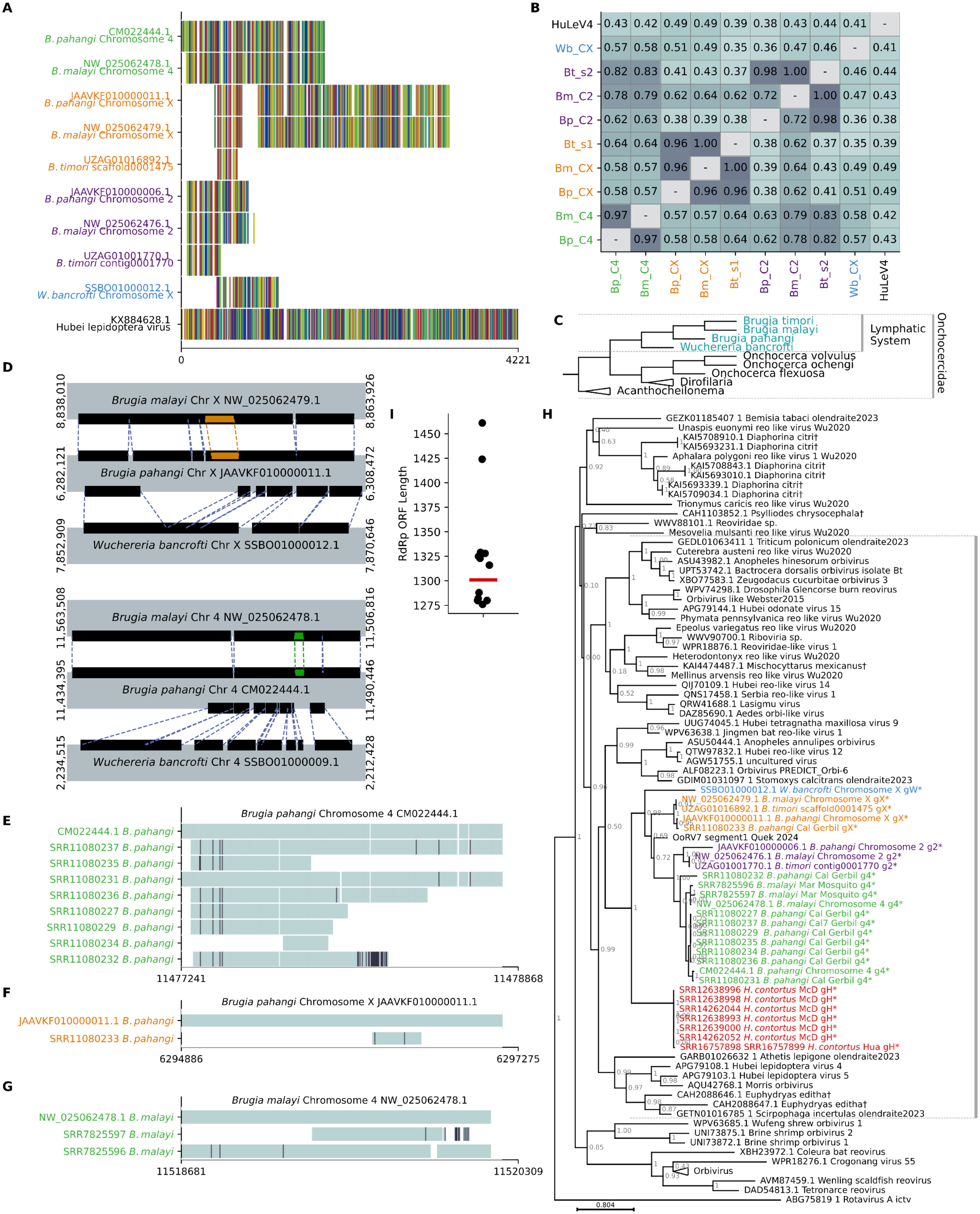
Orbi-like EVEs. (A) CIAlign [88] mini alignment showing the regions of the RdRp gene represented in chromosomal regions of *Brugia pahangi*, *Brugia malayi*, *Brugia timori* and *Wuchereria bancrofti* with similarity to orbivirus-like RdRp and their similarity to each other. Hubei lepidoptera virus 4 provides a full length reference RdRp. Text colours correspond to clusters based on genome position. Coloured lines represent different nucleotides (B) Similarity matrix showing the proportion of identical sites (excluding gaps) between the aligned EVE regions shown in (A). Abbreviations: *Brugia pahangi* Bp, *Brugia malayi* Bm, *Brugia timori* Bt, *Wuchereria bancrofti* Wb, chromosome C, Hubei lepidoptera virus 4 HuLeV4. (C) Simplified phylogenetic tree for the Onchocercidae, generated manually based on [61]. Clades for genera not discussed in the text are collapsed. (D) Schematic showing the regions surrounding the orbi-like EVE insertions on *Brugia* and *Wuchereria* chromosomes X (top) and 4 (bottom). Regions shown in black and linked by dotted lines were successfully aligned using NUCMER [90] Coloured regions represent the position of the EVE. (E-G) CIAlign mini alignments showing the sections of SRA assembled contigs which aligned to *B. pahangi* chromosome 4 (E), *B. pahangi* chromosome X (F) and *B. malayi* chromosome 4 (G). Teal regions are identical, white represents gaps and black represents differences from the chromosome. (H) Approximate maximum likelihood [84] phylogenetic tree for the newly identified orbi-like exogenous and endogenous viruses from nematode (labelled with *) and previously annotated sequences in this clade. Newly identified sequences are coloured by cluster and given a corresponding suffix - green, g4, chromosome 4-like; orange, gX, chromosome X-like; purple, g2, chromosome 2-like, red, gH, *Haemonconchus*-like; blue, gW, *Wuchereria*-like. Prefixes are GenBank or SRA accessions except for sequences with the suffix Wu2020, which are from [8]. The orbivirus clade has been collapsed and contains a monophyletic group which incorporates all ICTV classified members of the orbivirus genus. The clade marked with a grey line is the orbi-like clade referred to in the text (I) The distribution of ORF lengths of previously annotated RdRps in organisms in the marked clade (black dots) compared to those identified in the *H. contortus* contigs (pink line).

Using TBLASTX against the 229 available NCBI genomes (listed in Supplementary Data) from the phylum Nematoda, with known members of this clade as queries, fragments resembling the orbivirus-like RdRp were identified in two additional species, *Brugia timori* and *Wuchereria bancrofti*. These species are also members of the Onchocercidae and are the closest known relatives of *B. pahangi* and *B. malayi* [61,62], Figure 4A, C). They also both cause lymphatic filarial diseas and are vectored by mosquitoes. Together, these four species represent all known lymphatic filarial worms. Two distinct *B. timori* sequences were identified, one shorter than but highly similar to the chromosome X insertions in the other *Brugia* species (96-100% identity over 216 nt) and one to the chromosome 2 insertion (98-100% identity over 380 nt) (Figure 4A, B). The two *B. timori* sequences only overlap by approximately 50 nt. The *W. bancrofti* insertion is also short (664 nt) and is not very similar to any of *Brugia* insertions, with nucleotide identity from 35% to 58% (Figure 4A, B). The alignment on which these percentages are based is available in the Supplementary Data.

Given that the sequences on the same chromosomes in *B. pahangi* and *B. malayi* share high similarity, the positions of the sequences on the chromosomes were examined to establish if they share an integration site, and therefore are likely to have integrated into the genome prior to the divergence of the hosts from their last common ancestor. Based on chromosome to chromosome alignment between these two species, the chromosome 4 and chromosome X insertions are both at the same position in their respective genomes, within much longer homologous regions of the chromosomes (Figure 4D). The *B. pahangi* and *B. malayi* chromosomes align closely, giving strong evidence that these insertions occurred before the divergence of the two species. This kind of alignment was not possible for the chromosome 2 insertions, which may therefore be at different sites or within more divergent regions.

It was not possible to perform chromosome to chromosome alignment for *B. timori*, as in both cases the fragments were identified at the end of a short contig. However, after cropping the RdRp like region, the remainder of the contig which contains the chromosome 2-like insertion (UZAG01001770.1) shows 99% BLAST identity to the region of chromosome 2 which is adjacent to the insertion in *B. malayi*. A similar analysis with the chromosome X-like insertion did not identify a good alignment, however the *B. timori* contig is of low quality, with several long runs of unidentified bases. The *B. timori* genome is very fragmented (the scaffold N50 is 4.9 kb, compared to 10.9 Mb for *B. pahangi* and 14.2 Mb for *B. malayi*), which may explain the absence of an identifiable chromosome 4-like insertion in *B. timori*. Previously phylogenetic analysis has shown *B. timori* to share a more recent common ancestor with *B. malayi* than these two species share with *B. pahangi* [61,62], Figure 4C, the date of this ancestor is not established). If this is the case, the insertions which are at the same position in *B. malayi* and *B. pahangi* must also be at the same position in *B. timori*.

Integration prior to the divergence of these host species is consistent with the fact that the sequences cluster by chromosome rather than by host (Figure 4B). The sections of RdRp which are present in the three species also fit with this hypothesis - the chromosome 2-like and chromosome 4-like insertions all contain the 5’ region of RdRp (relative to Hubei lepidoptera virus 4) while the chromosome X-like insertion is more central (Figure 4A).

The *W. bancrofti* sequence is also on chromosome X but is divergent from the chromosome X insertions in the other three species. It was possible to align chromosomes X and 4 of *W. bancrofti* with those of *B. pahangi*, however the insertions did not occur within recognisable regions of similarity (Figure 4D). No sequence resembling RdRp was identified on *W. bancrofti* chromosome 4 but there is a gap in the alignment at the position containing the insertion in *B. pahangi*, so it is not clear if the absence is due to poor sequencing of this region. For chromosome X, the alignment is also fragmented, but the chromosome X region resembling RdRp in *W. bancrofti* (positions 20,061,483 to 20,062,162) is distant from the region which aligned to the areas surrounding the insertion in *B. pahangi* (positions 6,282,121 to 6,308,472). Given the low similarity between the *W. bancrofti* sequence and the *Brugia* sequences (Figure 4B) and the different region of RdRp which is represented (Figure 4A), it is likely to represent a different insertion event. If the insertions are not at the same positions in *W. bancrofti*, this means their insertion date was a maximum of approximately four to six million years ago, the estimated date of the last common ancestor of *Wuchereria* and *Brugia* [63].

To identify if any of these EVEs are transcribed, publicly available RNA-seq datasets from nematodes containing putative Reoviridae RdRp-like sequences were identified using the Serratus project database [11]. The datasets with putative Reoviridae were downloaded from SRA, filtered to remove reads mapping to the host and assembled into contigs, which were then screened using BLAST and HMMER to identify RdRp-like regions. Regions were identified in 18 datasets: 9 from *B. pahangi*, 2 from *B. malayi* and 7 from *Haemonchus contortus* (there are currently no publicly available RNA-seq datasets in SRA for either *B. timori* or *W. bancrofti*).

*H. contortus,* commonly known as the barber’s pole worm, is a comparatively distant relative of the lymphatic filarial nematodes, it is of the same order, Rhabditida, but it is not part of the filarial nematode (Filarioidea) superfamily. Instead, it is a pathogenic nematode of ruminants, causing the disease haemonchosis, which attaches primarily to the musoca of the abomasum (fourth stomach) [64]. Unlike the lymphatic filarial worms, it is not mosquito transmitted. The divergence between *Brugia* and the clade containing *Haemonchus* is estimated to have occurred 240 million years ago [65] and there are many species which are more closely related to *Brugia* which showed no evidence of reovirus-like transcripts or EVEs.

The *B. pahangi* and *B. malayi* sequences identified in SRA datasets do not contain intact open reading frames and instead resemble the endogenous insertions. All of the samples with reovirus-like regions in *B. pahangi* are from the same study (NCBI BioProject PRJNA606538), which appears to be currently unpublished, but involved experimental infection of Mongolian gerbils (*Meriones unguiculatus*) with *B. pahangi* and treatment with rifampicin of *Wolbachia*, the bacterial endosymbiont of *B. pahangi*. Nine of the twelve samples in this study expressed a reovirus-like transcript, from both treated and control gerbils and from both timepoints. Eight of the nine *B. pahangi* contig regions were highly similar to the chromosome 4 *B. pahangi* insertion and none extend into parts of the RdRp not included in this insertion (Figure 4E). The contigs containing the insertion are highly similar to each other and the RdRp represents the majority of the contig in all cases. It is likely that these transcripts are generated from the chromosome 4 EVE locus. The remaining contig region is identical to a short section of the chromosome X locus and is likely derived from this locus (Figure 4F). The two *B. malayi* contigs are from another seemingly unpublished study (NCBI BioProject PRJNA294263), in this case from experimentally infected *Aedes aegypti* mosquitoes. Both *B. malayi* samples from this study had reovirus-like reads. These two contigs appear to be derived from the chromosome 4 *B. malayi* locus, as again the ORFs are degraded and the region detectable in the contigs does not extend beyond the region encoded in the EVE (Figure 4G). Alignments are available in the Supplementary Data.

Phylogenetic analysis was performed incorporating the *H. contortus* sequences, reconstructed ORFs from the endogenous insertions in *B. malayi, B. pahangi, B. timori* and *W. bancrofti* and the related SRA sequences and orbivirus reference sequences (Figure 4H, newick file available in Supplementary Data). Historically, the RNA virome of nematodes has not been well known, however a number of new viruses were identified in a recent virus discovery project [66]. This publication identified two Reovirales-like sequences in nematodes, one in another member of the Onchocercidae family, *Onchocerca ochengi*, a filarial (but not lymphatic) nematode of cattle closely related to *Brugia* and one in *Haemoconchus contortus*. These were incorporated into the phylogeny. Our nematode sequences fall within a single clade with branch support of 96%, together with the *O. ochengi* sequence, but surprisingly not the *H. contortus* sequence. The *H. contortus* sequence from this recent publication did not cluster with the orbi-like viruses in our analysis but was identified in the same BioProject as some of our *H. contortus* RdRps.

As expected from the results above, the *Brugia* sequences cluster phylogenetically by chromosome rather than by host, with distinct, well-supported chromosome 2, chromosome 4 and chromosome X clades. The chromosome 2 and chromosome 4 insertions and their associated SRA transcripts cluster closely together, while the chromosome X insertions are somewhat more distant. The *H. contortus* sequences form a separate, monophyletic group to the viruses of filarial nematodes. The phylogenetic relationships in this tree suggest that the nematode orbi-like viruses share a common ancestor more recently with each other than they do with the insect orbi-like viruses. However, the virus phylogenetic tree reflects several independent endogenisation events.

Unlike the *Brugia* contigs, the contigs derived from RNA-seq in *H. contortus* show hallmarks of being an active, circulating viruses. The genome of *H. contortus* has been sequenced and assembled at chromosome level (GCA_000469685.2) but does not contain any regions resembling reoviral RdRp, including in a second screen using the assembled *H. contortus* sequences as queries. Four of the SRA assembled contigs have a complete, 1,301 amino acid open reading frame, comparable to exogenous viruses from this clade (Figure 4I, Supplementary Data). It would usually not be evolutionarily advantageous for an EVE to maintain an ORF of this length, capable of producing intact viral proteins, for any period of time. For the remaining three sequences the ORF is truncated at the 3’ end, likely because the contig assembly is incomplete. The ORF makes up 96-98% of the full contig length, again consistent with known, related viruses. These results are consistent with transcription from an intact, circulating RNA virus, rather than an EVE.

Six of the seven *H. contortus* datasets containing reovirus-like contigs are from BioProject PRJNA264197 [67], covering six of 27 laboratory passaged *H. contortus* samples subjected to various treatments with small molecule inhibitors. The other dataset is from BioProject PRJNA756739 [68], in one of seven samples from experimentally inoculated goats. Therefore, the presence of this virus in wild populations of *H. contortus* cannot be confirmed.

Interestingly, the recently identified sequence from *O. ochengi* [66] also resembles an intact, exogenous virus. It has an 1,299 amino acid RdRp open reading frame covering a large majority of the contig. The reference genome of this species, assembled to scaffold level, was screened for endogenous insertions but did not appear to share any regions of homology with the orbi-like viruses. This host species is much more closely related to *Brugia* than *H. contortus* (Figure 4C), and the virus sequence is more similar to that of the *Brugia* EVEs.

If the virus is exogenous in *H. contortus*, it is expected that sequences resembling the other orbivirus segments besides RdRp will be present in these datasets. Members of the Orbivirus genus have 10 segments and encode 12 proteins [69]. However, some segments have never been identified for members of the phylogenetic clade containing the nematode viruses (marked on Figure 4H). Between one and eight sequenced proteins are available from NCBI for viruses in this clade, these were compared with Pfam profiles for ten orbivirus proteins (VP1 to VP7 and NS1 to NS3). A maximum of four proteins could be identified for any single virus using this approach, and all identified proteins matched VP1, VP3, VP4 or NS1. Wu et al 2020 [8] also sequenced a number of viruses in this clade but, unlike for other reoviruses in their study, did not find any additional segments, suggesting segments in this clade are divergent.

Using a combination of HHsearch, HMMER, BLAST and predicted structure comparison with FoldSeek, VP1 (RdRp), VP3, VP4 could be identified amongst *H. contortus* contigs in all seven SRA samples. FASTA files containing contig sequences and ORFs are available in the Supplementary Data. Phylogenetic trees based on VP3 and VP4 were not substantially different from the tree based on VP1 (Figure 4H, Supplementary Figure 9). The VP3 and VP4 ORFs were consistent with the length of those identified for other members of the clade (Supplementary Figure 9).

Unidentified proteins from viruses in this clade were also compared to the *H. contortus* contigs, however no convincing similarity was found. Segments from members of the orbivirus genus are known to have terminal repeat sequences, usually six nt in length, which are somewhat conserved within species [70], however these could not be found either for the *H. contortus* contigs or for any previously identified members of this clade.

### Virus Labelled Sequences

Besides the EVE sequences, 1,609 uncharacterised proteins were analysed which are already classified as viral in NCBI but not specifically as either Orthornavirae or RdRp. For 1,541 sequences the lowest taxonomic classification is the realm “Riboviria” (NCBI Taxonomy ID 2559587), which includes the Orthornavirae kingdom but also the Pararnavirae, which encode RT rather than RdRp, and several groups of satellite viruses (Figure 5A). The remaining 68 sequences are only classified at the superkingdom level, as “Viruses” (NCBI Taxonomy ID 10239). Classifications are provided in Supplementary Table 1. These sequences are RdRp proteins from RNA viruses which either have not been officially taxonomically classified, or which were yet to be classified at the time of their submission to the database. They are not labelled as RdRp but as uncharacterised proteins, however, all meet the criteria used here to classify a protein as a full or partial RNA viral RdRp.

**Figure 5:**
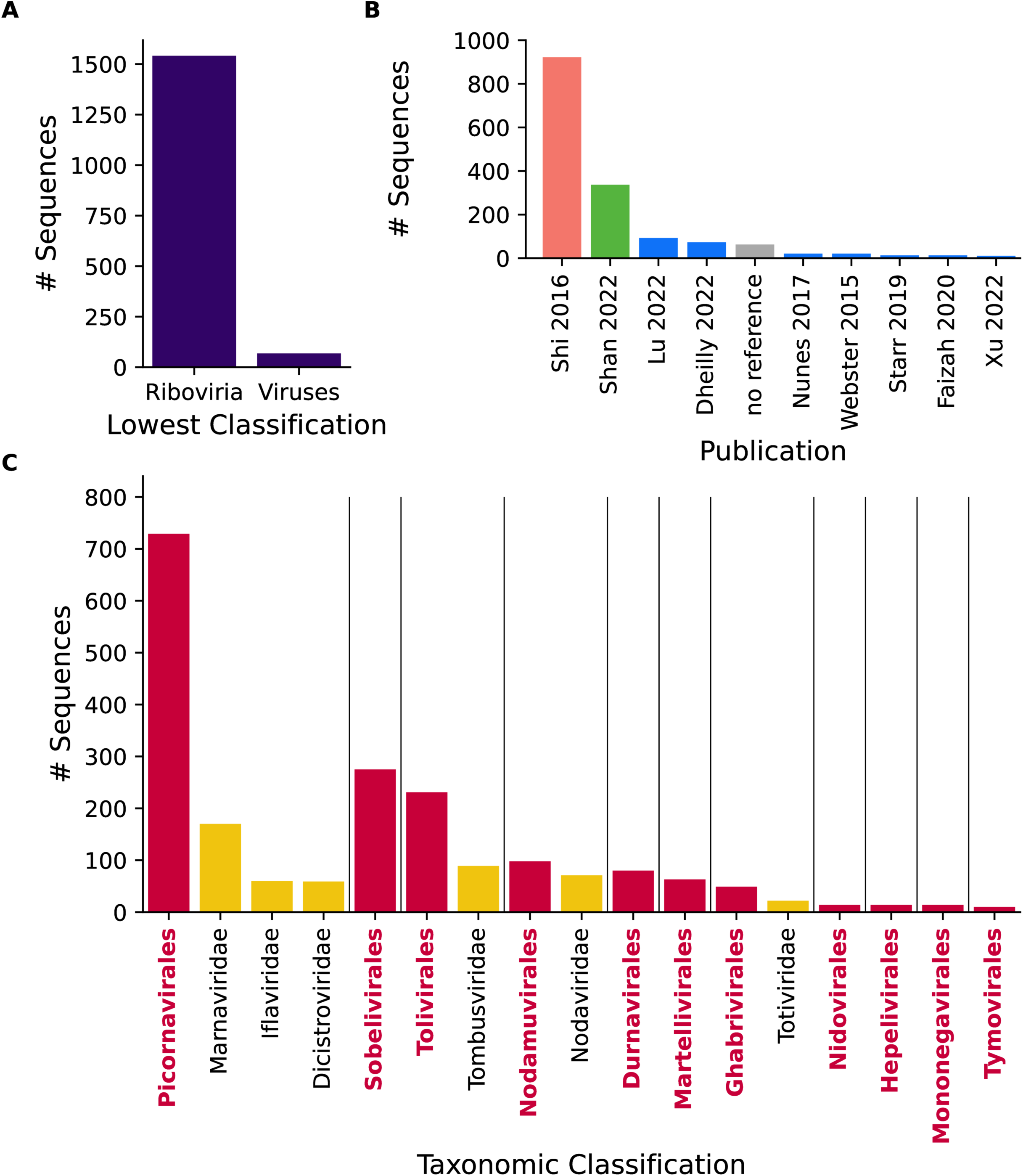
Virus Labelled Sequences. (A) Bar chart showing the number of uncharacterised proteins with a previous lowest taxonomic classification of Riboviria or Viruses. (B) Bar chart showing the number of unchacterised RdRp-like proteins from specific publications, labels correspond to publications as follows: Shi 2016 [2], Shan 2022 [71], Lu 2022 [91], Dheilly 2022 [92], Nunes 2017 [93], Webster 2015 [94], Starr 2019 [95], Faizah 2020 [96], Xu 2022 [97]. The “no reference” records were not associated with a specific PubMed ID. (C) Bar chart showing the ICTV RNA virus orders (pink, bold) and families (yellow) to which the virus labelled uncharacterised proteins could be assigned. Each section shows families within the named order.

The majority of the unassigned viral proteins come from a few specific publications (Figure 5B). Of the 1,609 sequences, 922 are from Shi et al’s 2016 publication [2], which described many novel groups of invertebrate viruses, the majority of which are yet to be officially classified by ICTV. 337 are from Shan et al. 2022 [71] who identified RNA viruses in the cloaca of many bird species. Of the remainder, 287 are distributed across 26 other virus discovery publications and 63 are not linked to a specific publication (Supplementary Table 1, Figure 5B).

Where possible, these sequences were provisionally assigned here to an ICTV classified family. Only sequences which fell within the existing diversity of a well supported (branch support ≥ 0.8), monophyletic cluster of ICTV classified viruses from the same family were assigned to that family. This is a conservative approach which may cause more divergent members of the same family to be excluded. 510 viruses were classified at family level using this approach. The largest groups were the Marnaviridae, with 170 members, Tombusviridae with 89 and Nodaviridae with 71 (Figure 5C). 1,604 sequences were putatively assigned to viral orders, most commonly the Picornavirales (729 sequences), Sobelivirales (275 sequences) or Tolivirales (231 sequences) (Figure 5C).

### Sequences Misclassified as Bacterial

251 RdRp-like sequences were identified amongst sequences classified as bacterial (Figure 1B). Of these, 96 are almost identical sequences from DNA contigs assembled from the *Mesorhizobium* species of nitrogen-fixing bacteria (Supplementary Data: Tree Cryppavirales_11). These sequences are all from the same study into root nodules from chickpeas (*Cicer arietinum*) and have a highly significant TBLASTN hit (99% identity across 100% of the protein) against chickpea chromosome 7 (AHII03000007.1), so it is very likely that these represent an EVE from the host of the bacteria which as been misclassified.

Of the remaining RdRp-like unclassified sequences labelled as bacterial DNA, unexpectedly, 56 sequences had very high BLASTP similarity scores (>85% identity and bit score > 400) against diverse ICTV classified virus proteins (Figure 6A, Supplementary Table 1). Bacteria are not known to have EVEs and it seems unlikely that so many bacteria, from multiple bacteria, would match well-known viruses so closely without this being previously detected. Therefore, these sequences were examined in detail.

**Figure 6:**
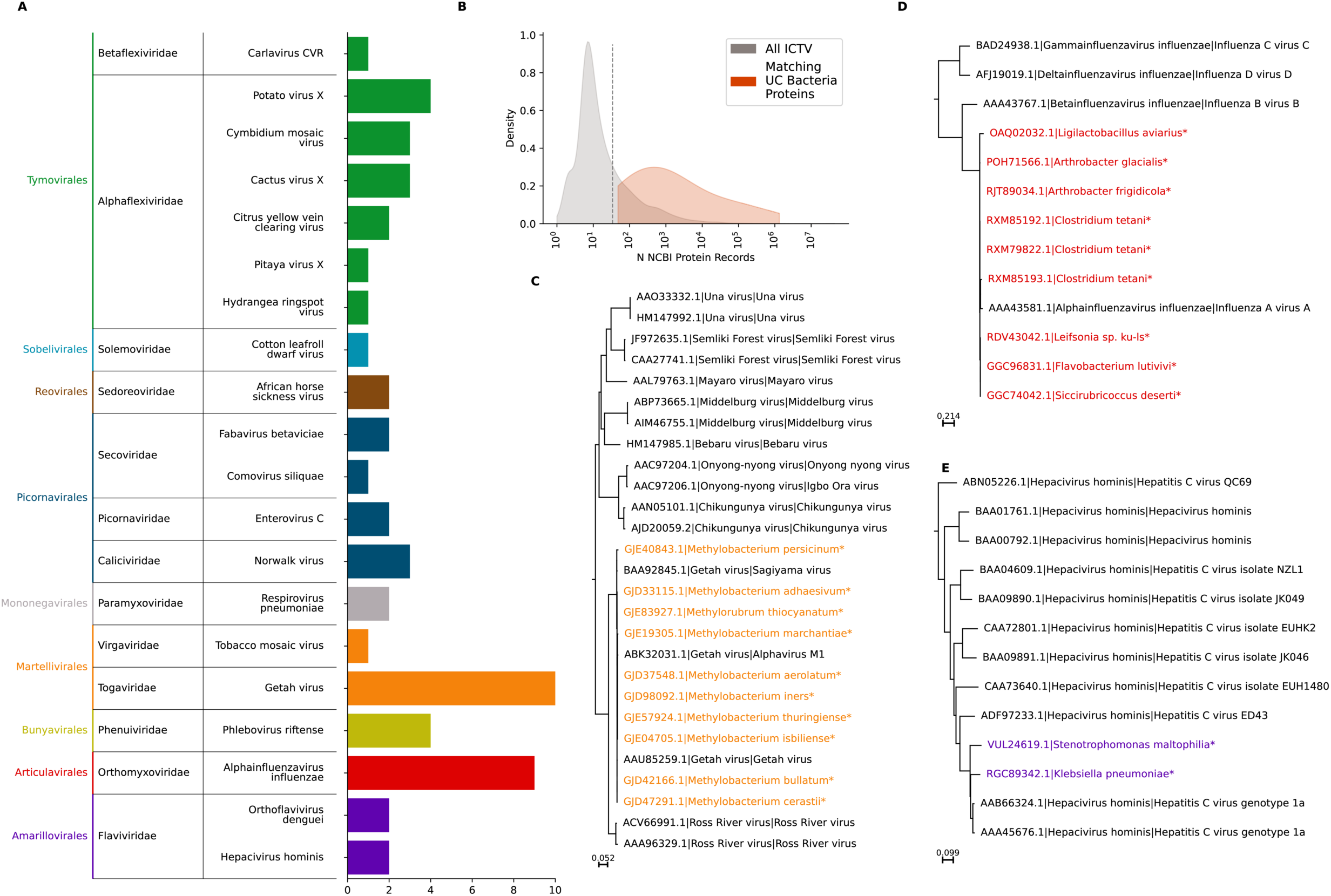
Sequences Mislabelled as Bacterial. (A) Bar chart showing, for sequences labelled in their original database as bacterial sharing 80% BLAST identity and a bit score >400 with an ICTV classified virus, the number of sequences resembling each virus. Colours and labels in the left column represent viral orders. (B) Density plot comparing the number of GenBank records per species for all ICTV classified viruses (grey) and for those which matched a bacterial-labelled RdRp-like uncharacterised sequence (red). (C-E) Approximate maximum likelihood [84] phylogenies for specific examples of bacterial derived sequences and previously identified sequences in the same clade, for sequences related to Getah Virus (C), influenza virus A (D) and Hepatitis C virus (E). Previously annotated sequences are labelled with their GenBank accession, then ICTV species name, then common name.

All of the 56 sequences were found to match very commonly studied pathogenic viruses, including widespread pathogens of humans and agriculturally important animals and plants, such as the viruses causing influenza A, dengue fever, hepatitis C and African horse sickness (Figure 6A). As a proxy for how widely studied a virus is, the number of NCBI protein records was calculated for all ICTV-classified RNA viruses and compared to the number for ICTV-classified RNA viruses with matching bacterial proteins (Figure 6B). ICTV-classified RNA viruses have a median of 10 protein sequences available, while those with matching bacterial proteins have a median of 778. This supports a hypothesis that the sequences labelled as bacterial are the result of either mislabelling or contamination with common laboratory strains.

Phylogenetic analysis further supported this hypothesis, with the bacteria-labelled sequences indistinguishable from known pathogens. For example, “bacterial” sequences were identified which fell well within the known diversity of Getah virus (Figure 6C), Alphainfluenza virus A (Figure 6D) and Hepacivirus homininis (Figure 6E). EVEs would be expected to have diverged from the original sequence somewhat, especially in the case of these highly pathogenic viruses. Tree files are available in the Supplementary Data.

These results together suggest that these are likely sequences from exogenous viruses which are the result of contamination of bacterial samples with viral RNA or mislabelling of RNA viral samples as bacterial.

### CDD Annotations

Sequences uploaded to the NCBI Protein database are automatically annotated via RPS-BLAST with conserved protein domain footprints from the conserved Domain Database (CDD) [72]. The CDD includes domains from RNA viral genes and therefore has potential in identifying RdRp-like regions amongst unidentified sequences. The CDD annotations of the uncharacterised RdRps identified here were therefore examined to assess whether the utility of these annotations in identifying viral proteins (Figure 7). CDD annotations were divided into four categories - specific RdRp domains (RdRp), explicitly RNA viral but not RdRp specific domains (RNA viral non-RdRp), reverse transcriptase-like domains (RT-like) and domains which are not specific to viruses (Other). The CDD annotations for each RdRp-like protein are provided in Supplementary Table 1.

**Figure 7:**
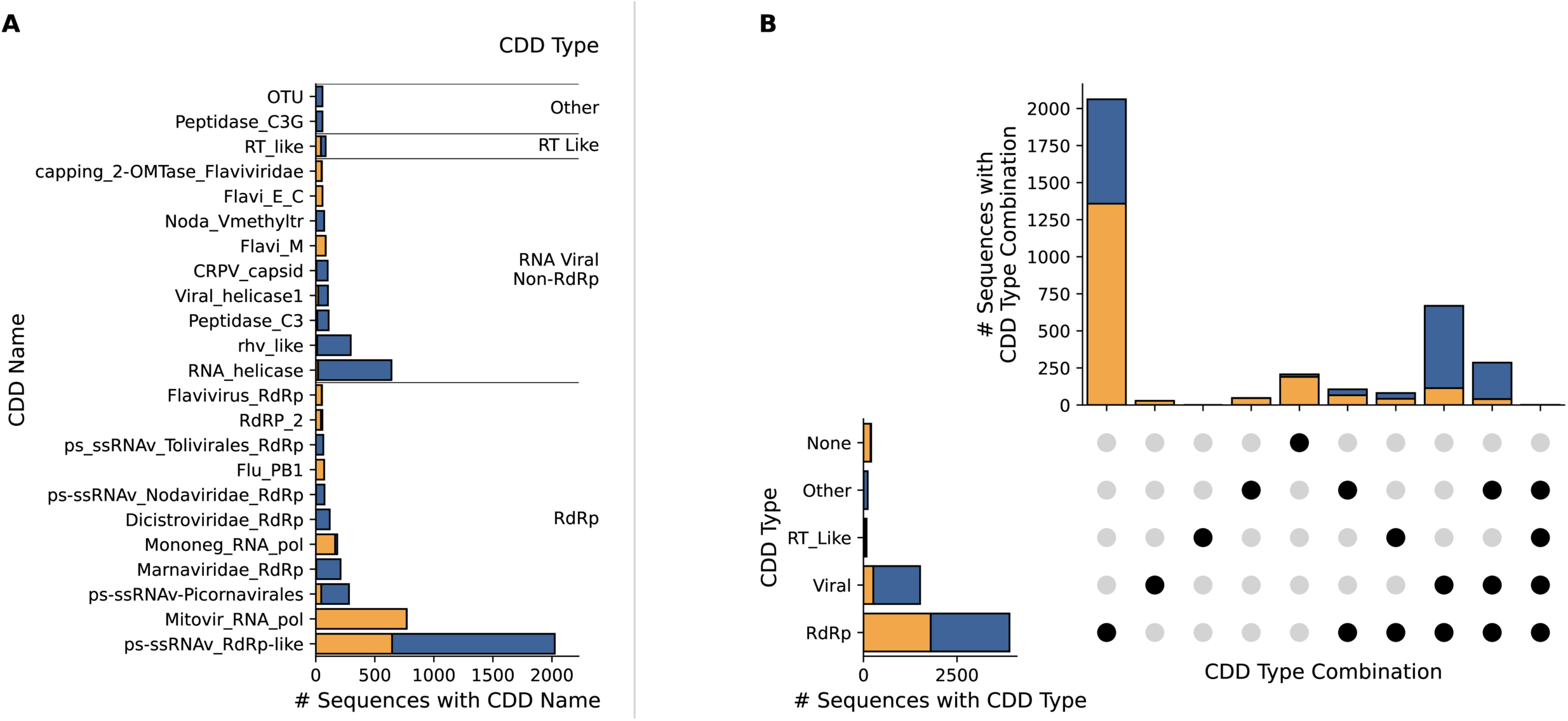
Conserved Domain Database Annotations. (A) Bar chart showing the proportion of RdRp-like uncharacterised proteins with CDD [72] annotations classified as RdRp-like, RNA viral non-RdRp, reverse transcriptase (RT)-like or other (not specifically RNA viral). Sequences not previously labelled as viral are shown in orange and on the left, those previously labelled as viral in blue and on the right. Only domains with >50 annotations in our dataset are shown (B) UpSet plot [98] showing the number of uncharacterised proteins with each combination of CDD annotations.

For the sequences previously assigned as “Viruses” or “Riboviria”, the CDD annotations provide a good estimation of whether a sequence is RNA viral (Figure 7A). Of 1,605 sequences, 1,585 have an RdRp-like domain, most commonly “ps-ssRNAv_RdRp-like”, the conserved catalytic core domain of positive stranded RNA viral RdRp (781 sequences) and ps-ssRNAv-Picornavirales, the Picornavirales specific version of this domain (281 sequences). This suggests that the CDD annotations are a useful tool in assigning an unclassified viral protein as RdRp. 20 RdRps labelled as viral were not annotated with RdRp and 17 had no annotations at all. All of these 17 sequences have a BLAST bit score > 100 against another viral RdRp and all were classified as RdRp by at least two r methods. The virus classified sequences often also had a CDD annotation of a second viral protein, most commonly helicase, which was rare for the sequences not originally labelled as viral (Figure 7A, 7B). This probably reflects the frequency with which these sequences represent exogenous, intact viruses rather than integrated EVEs.

The sequences previously assigned as non-viral also usually had a CDD annotation - 1,622 of 1,889 NCBI protein results not classified as viral had an annotated CDD RdRp (Figure 7A). The most common was Mitovir_RNA_pol (770 proteins), representing mitoviral RdRp and reflecting the high number of Mitoviridae EVEs identified in this group. Most of the non-viral sequences were only annotated with a single RdRp (Figure 7B).

These results suggest that CDD annotations are a useful tool in determining if a sequence is potentially RNA viral in origin, but they are not adequate without additional analysis. They are also not always easily accessible or interpretable - while the domain ID is provided in the NCBI record for the protein, it is not possible to search by domain and taxonomic information and other metadata about the domains is spread over a number of source databases with different interfaces and APIs.

### Near Misses

Another aim of this study was to identify proteins which are not RdRp but which score highly in HMM-based comparisons with RNA viral RdRp. To this end, over 1 million proteins were examined which had a HMMER score against a viral RdRp profile which was ≥10 and ≤25, described hereafter as “near miss” proteins. These were compared to a randomly selected control group of uncharacterised proteins for which HMMER did not return a score against RdRp. The likely functions of both groups of proteins were determined using BLASTP against proteins of known function and terms associated with these functions were categorised into groups. 577,704 near miss proteins had an identifiable probable function, and these were matched with 499,240 identifiable proteins from the control group. Figure 8A shows the 10 terms which were most significantly overrepresented amongst the near miss proteins compared to the control (also listed in the Supplementary Data). A list of the uncharacterised near miss proteins and their BLAST target sequences is provided in the Supplementary Data.

**Figure 8:**
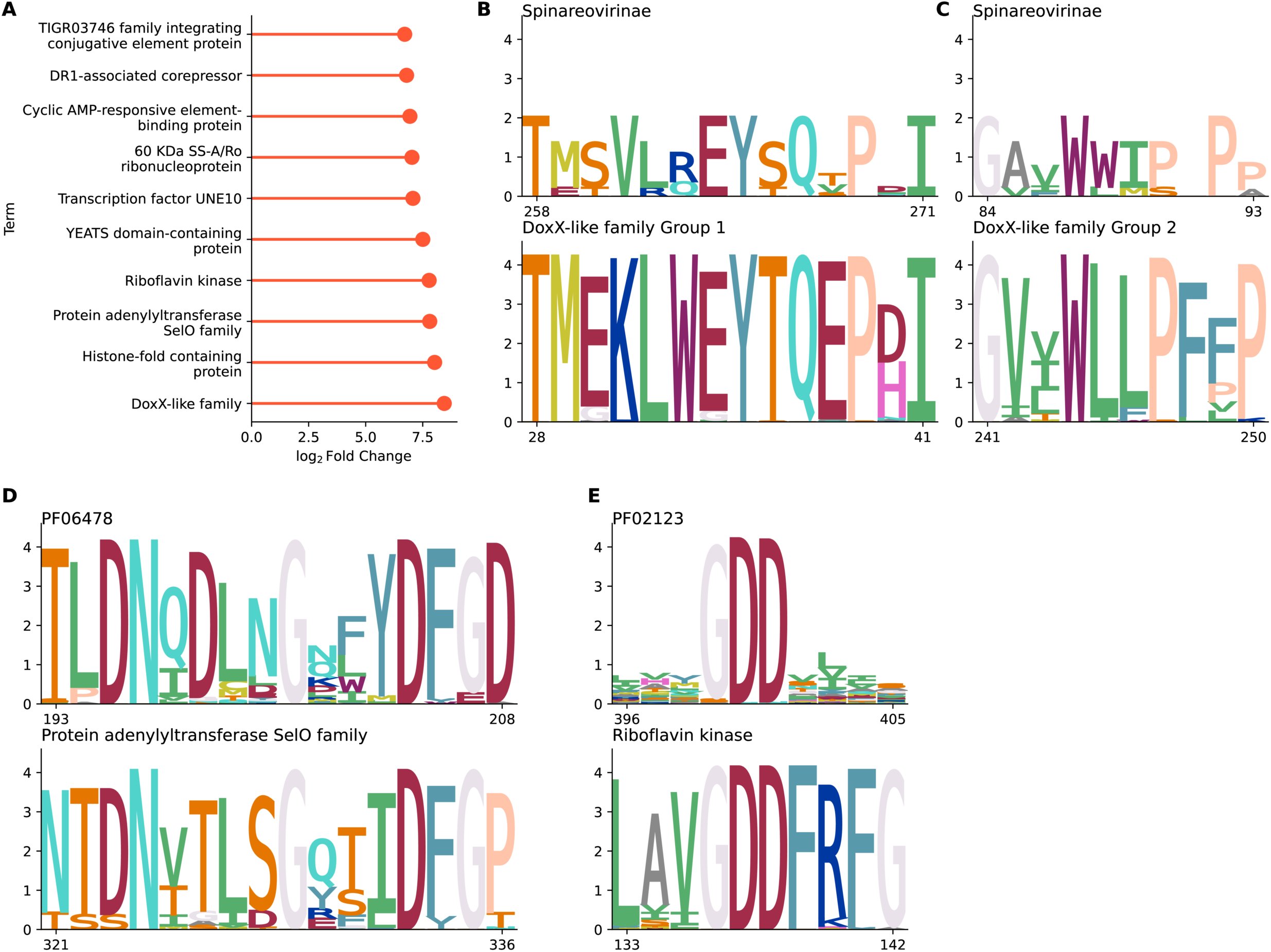
Near-miss Analysis. (A) Bar chart showing the ten terms which were most overrepresented amongst BLAST hits against uncharacterised proteins scoring between 10 and 25 in HMMER [81] comparisons with viral RdRp pHMMs compared to uncharacterised proteins with no score. Bars represent the log_2_ fold change between the frequency of the term in the two sets. (B-E) Sequence logos generated with CIAlign [88] showing the alignment between specific regions of the near miss proteins and the pHMM profiles against which they had a weak match, for DoxX-like family proteins (B, C), protein adenylyltransferase SelO family proteins (D) and riboflavin kinase proteins (E). Positions in the near miss proteins are relative to the alignments provided in the Supplementary Data, positions in the profiles are relative to the original profile.

Proteins related members of the DoxX-like family of integral membrane proteins were particularly overrepresented, 785 of the near miss proteins were related to proteins annotated with this term, compared to two of the control proteins. The BLAST target sequences most similar to the uncharacterised proteins were 191 closely related proteins approximately 300 amino acids in length, from the *Bacillus* genus of bacteria. An MSA for these sequences is available in the Supplementary Data. Two distinct regions were identified resembling the HMM profile for the Spinoreovirinae RdRp, labelled here as group 1 and group 2 for clarity (Figure 8B). The group 1 domain (positions 23 to 76 in the MSA) has similarity to part of the StAR-Related Lipid-Transfer (START) domain as well as to Reoviral RdRp, while the group 2 region (positions 239 to 271 in the MSA) does not match any specific domain of the DoxX-like proteins. Lipid transfer proteins do play a role in the life cycle of some RNA viruses [73] and it is interesting that both regions match specifically the Spinoreovirinae, but there is not enough evidence here to infer anything other than coincidental similarity. Nevertheless, DoxX-like proteins could be incorporated into negative control protein sets to test the specificity of RdRp-detection methodologies.

A moderate level of similarity to the Pfam profile for Coronavirus RNA-dependent RNA polymerase, N-terminal (PF06478) was identified amongst unidentified proteins with similarity to 56 bacterial proteins, mostly from the genus *Corynebacterium*, annotated as protein adenylyltransferase SelO family proteins (MSA provided in Supplementary Data, high similarity was detected at positions 297 to 348). These are pseudokinase proteins involved in cellular response to oxidative stress [74]. Again, there is no particular reason to suspect these proteins are related, but there is visible similarity between the coronavirus RdRp and the SelO proteins within a specific region (Figure 8C).

A set of riboflavin kinase genes from various bacterial genomes also show some similarity to the near-miss proteins (MSA provided in Supplementary Data, high similarity was detected at positions 132 to 168). In this case, the conserved GDD motif which is common in RNA viral RdRp [75] is apparent in the bacterial alignment

## Discussion

By screening only the uncharacterised proteins in these two large databases for a single viral gene, a surprising number of RNA viral RdRp proteins, particularly RNA viral EVEs integrated into eukaryotic genomes, were identified. These results both reveal the extensive diversity of unknown EVEs and how they can impact our understanding of contemporary viruses and demonstrate the complexities of generating a negative control datasets for RNA virus discovery.

It is clear from our results that non-RNA viral proteins from the NCBI protein or UniProt databases, used without additional filtering, are not a good negative control in RNA virus discovery research, as they contain “false false positives” - sequences which will show as false positives (RdRps in the negative control) but are actually mislabelled true positives. The approximately 3,500 proteins described here were all identified as RNA viral RdRp via at least two sources of evidence and fall comfortably within RNA viral phylogenies. It is therefore recommended that, if these databases are used to eliminate potential false positives, uncharacterised proteins are first removed. Additional putative false positives against named proteins should be carefully checked to ensure that the named proteins are not from an RNA viral or EVE source.

Besides their utility in future methods development, the results presented here also show that EVEs remain under-studied and under-characterised. This study had a relatively limited scale compared to much virus discovery research, but uncovered hundreds of previously unknown elements. Plants and insects in particular were, as expected, a rich source of EVEs from diverse virus families. EVEs were also found in unexpected taxa, for example the Perkinsozoa phylum of protists and the parasitic worm *Pomphorhynchus laevis*, for which almost nothing is currently known about the RNA virome.

Our findings for the Cryppavirales suggest recent or ongoing transfer of genetic material from mitoviruses to plant genomes. Our results are consistent with the currently accepted hypothesis that plant mitoviral EVEs originated from fungi [76–78], in that our plant mitoviruses fall almost exclusively in the monophyletic Duamitovirus genus. A single clade within this genus which contains almost exclusively plant endogenous and exogenous viruses, distinct from the fungal members of the genus. Members of this clade have specific RdRp features which are not apparent in other mitoviruses, potentially suggesting some degree of adaptation to plant hosts. Mitoviruses in plants are considered to have initially integrated in the mitochondria and later been passed to the nuclear genome by DNA transfer [76]. Around one in four of our sequences were identified in mitochondria, but many additional sequences are from unassigned contigs, so a mitochondrial origin cannot be ruled out. However, where contigs were assigned to chromosomes, identical or near identical mitochondrial and nuclear sequences were common, for example for the 100% identical Capiscum sequences shown in Figure 1H. This suggests that, at least in these cases, if the nuclear EVEs are integrated mitochondrial DNA then the integration has occurred recently, which is feasible since the process of mitochondrial transfer of DNA into the nucleus is considered to be ongoing in plants [79]. Older insertions into the nuclear genome would be expected to have degraded to some extent. Further evidence for the recent integration of many of the mitoviral EVEs identified here comes from the lack of congruence between the host and EVE phylogenies, with little evidence of insertions coevolving with their hosts. Based on our results, integration of mitoviral EVEs seems to be recent, diverse and widespread across at least the Magnoliopsida class of flowering plants.

Other EVEs identified here appear to be much more ancient. For example, the large cluster of fungal viruses related to the Curvulaviridae (Figure 3A) contains likely exclusively endogenous sequences with which the host phylogeny can be almost fully recapitulated. It is likely that the viruses found here are older than the divergence of this group from a common ancestor and that if further members of this family were subjected to genomic sequencing, more examples would be identified.

The results for the orbi-like viruses present an interesting case, with exogenous and endogenous viruses in specific, distantly and closely related, nematode species. For the *Brugia* orbi-like viruses, there is no evidence of ongoing exogenous viral activity, however there have been at least three separate integration events, seemingly prior to the divergence of the species of *Brugia* from their common ancestor. The EVEs in the genome appear to be ancient and are moderately degraded. However, the virus identified in *O. ochengi* [66] is a close relative of the endogenous insertions in *Brugia* and shows hallmarks of being a currently circulating, exogenous virus, suggesting ongoing activity by viruses in this clade throughout its evolutionary history. Meanwhile, the *H. contortus* sequences, also apparently exogenous, form a separate monophyletic group. The fact that these closely related viruses were identified in both *Haemonconchus* and the filarial nematodes, despite their large genetic distance, implies both that there are likely additional orbi-like viruses amongst the nematodes still to be discovered and that nematodes may be prolific hosts for members of this clade.

In conclusion, RdRp-like sequences, derived from both exogenous viruses and EVEs, are common amongst the uncharacterised proteins in public protein databases. Many EVE proteins in particular remain uncharacterised and provide a rich, largely untapped, source of data about virus host range and evolutionary history. RNA virus detection studies are becoming more common and in many cases reveal thousands or hundreds of thousands of previously unknown RdRps, but detailed characterisation of the RNA viral diversity amongst a set of sequences still reveal a great deal of novel insights.

## Materials and Methods

To identify uncharacterised and poorly characterised proteins, six search terms were used: uncharacterized, unclassified, unknown, unnamed, hypothetical, DUF. These are referred to henceforth as the “unknown protein search terms”.

Proteins were included in the target database as follows: National Centre for Biotechnology Information (NCBI) Protein database [21] (accessed 2022-08-08) proteins containing any of the unknown protein search terms in their “title” field and not classified taxonomically as “Orthornavirae” and proteins from UniRef100 [25], accessed 2023-05-31, 2023-05-03 EBI release) which contain any of the unknown protein search terms or ‘putative’ in the protein name.

Three sets of publicly available pHMMs were selected to allow for comprehensive screening. Pfam profiles are based on alignments extracted from the HHsuite-3 pfamA 35.0 database [80] (accessed 2023-02-17, 2021-11-23 release), for 22 Pfam families identified as RNA viral RdRp, these are listed in the Supplementary Data. TSA profiles are the 74 pHMMs described by Olendraite et al., 2023 [16]. RdRp-Scan profiles are the 43 pHMMs described by Charon et al., 2022 [26]. All pHMMs were created from the provided multiple sequence alignments using HMMbuild from HMMER v3.3 [81].

pHMMs were compared with the target database using HMMscan from HMMER, with the default settings. HMMER results were filtered to retain only sequences with a domain score ≥ 25. For the near miss analysis results were filtered to retain sequences with a domain score ≥10 and ≤25. RefSeq records were removed if they were copies of another sequence already in the database. Sequences present in the RdRp-scan database [26] and therefore already known to be RdRp, were also excluded.

Details of filtering reverse transcriptase hits from Pfam profiles PF05919 and PF00680 are provided in the Supplementary Materials and Methods.

### Virus BLAST Searches

DIAMOND BLASTP (v0.9.14) [28] against the non-redundant BLAST protein database (nr, downloaded 2023-06-14) was used to further interrogate the RdRp-like proteins. BLASTP results were filtered to exclude self hits. The most similar ten target proteins not labelled as uncharacterised were identified. If there were no matches against characterised proteins, the search was repeated using BLASTP via the online NCBI BLAST server (https://blast.ncbi.nlm.nih.gov/Blast.cgi, accessed 2023-08-08) and the results processed as for the DIAMOND BLAST results. Sequences matching only RdRp were classified as “RdRp”, those matching RdRp and another adjacent named protein as “RdRp plus adjacent”, those matching reverse transcriptase, maturase or integrase as “RT-like” and those matching only a named protein which is neither RdRp or RT as false positives. Target proteins were examined manually to ensure they were assigned to the correct category. Proteins classified as RNA virus or RNA virus plus adjacent were considered to be true positives, all others to be false positives. Proteins matching only uncharacterised proteins were retained but required a further line of evidence (as described below) to be classified as RdRp. Three named proteins in nr were found to be likely mislabelled RNA viral proteins, these were ACF19853.1 (ANT-5, *Toxicara canis*), KMQ93496.1 (gag-pol fusion protein, *Lasius niger* and KMQ91513.1 (glucuronate isomerase, *Lasius niger*), therefore matches against these three sequences were considered to be true positives. The best BLASTP match (excluding uncharacterised sequences), BLAST methodology used (DIAMOND BLAST or online BLASTP) and BLAST bit score against this match are listed for all proteins in Supplementary Table 1.

### Clustering

Proteins passing this initial filtering were clustered using the MMseqs2 (v14.7e284) [82] cluster algorithm at 85% similarity and 25% coverage in cluster mode 0 and with ‘--cluster-reassign’ enabled. The representative sequences were aligned to the sequences making up the HMM profile to which they had the most significant match and provisional phylogenetic trees were generated, using MAFFT (v7.520) [83], and then FastTree (v2.1.11) [84], both with the default settings. Based on their clustering in this analysis, sequences were assigned to one of 32 virus groups based on the profile to which they matched. These groups are predominantly at order level, with the exceptions of the phylum Lenarviricota (for which many published sequences were not classified at a lower level), the families Birnaviridae and Permutotetraviridae, which do not have a higher level International Committee on Virus Taxonomy (ICTV) classification and the provisional genera Zhaovirus, Yanvirus and Weivirus [2] and Quenyavirus [85], which have not yet been classified by the ICTV.

Within these order-level groups, existing viral sequences have been subdivided, based on our previous phylogenetic and clustering analyses, into 331 operational taxonomic units (OTUs). Clusters of uncharacterised proteins were expanded into the original sequences and each sequence was sequentially incorporated into the phylogenetic tree for each OTU within its assigned order-level group, using MAFFT and FastTree settings as above. Sequences were assigned to the OTU within which they had the shortest distance (calculated from the trees with the ete3 v3.1.2 [86] get_distance function) to any known virus leaf. The OTU assigned to each sequence and its distance from the nearest neighbour within this tree are provided in Supplementary Table 1.

### Filtering HMMER Results

To increase the probability that identified sequences are truly RNA viral, additional screening was performed with Foldseek, PalmScan and BLAST and using the phylogenies generated above.

Sequences were classified as likely RNA viral if they met two of more of the following criteria (software details below): 1) A high FoldSeek bit score of >100 against an RdRp profile, 2) an identified RdRp core using PalmScan, 3) a high BLAST bit score of >50 against a known RdRp sequence in the BLASTP analysis described above, 4) a branch length of less than 1.5 separating the sequence from the nearest known RNA virus in phylogenetic analysis or 5) a HMMER score >50 against an RdRp profile.

Foldseek analysis was performed to identify proteins with protein structural similarity to RdRp. Putative RNA viral proteins were split into substrings with a length of 400 amino acids and an overlap of 200 amino acids (as this is the maximum size which can be processed). These sequences were folded using the ESM Fold Sequence [87] online server (accessed via API on 2023-07-04 at https://api.esmatlas.com/foldSequence/v1). Similarity scores were then calculated using FoldSeek v6.29e2557 [29] against 141 PDB profiles identified as RdRp (listed in Supplementary Data).

PalmScan v1.0.i86linux64 [18] was used to identify core RdRp motifs in all proteins, with RdRp and RT detection enabled and otherwise the default settings.

Phylogenetic tree distances were calculated with the ete3 get_distance function as above. HMMER and BLAST scores were calculated as discussed above.

### Phylogenetics

Phylogenetic trees for each viral OTU were regenerated containing only the sequences passing these filtering criteria. To generate these trees, sequences were aligned with MAFFT under the default settings as above, then alignments were cleaned with CIAlign (v1.1.4) [88] to remove insertions up to 1000 amino acids in length, remove terminal regions with coverage <10% or similarity <10% and remove sequences <50 amino acids in length. Trees were then generated with FastTree as above. These trees are available in the Supplementary Data.

Unless otherwise specified, subtrees in figures were generated by manually selecting clades from these large phylogenies, realigning sequences at higher stringency using the MAFFT linsi algorithm (v7.520) [83], cropping with the CIAlign to remove terminal regions with coverage <50% or similarity <30% and to remove sequences <50 amino acids in length, and rebuilding trees using FastTree as above, with additional reference sequences added in some cases. Subtrees were visualised using plot_phylo (v0.0.4, <u>github.com/KatyBrown/plot_phylo</u>).

Methodology used specifically to characterise the Cryppavirales and Reovirales are provided in the Supplementary Materials and Methods.

### Virus Derived Sequences

To classify proteins already labelled as viral, a BLASTP search (v2.14.0) [30,31] was performed for uncharacterised proteins from records labelled as “Riboviria” or “Viruses”. These were compared to RdRp proteins linked from the nucleotide accessions listed in the ICTV virus metadata resource (version VMR_19-250422_MSL37) from 5,000 viruses across 3,905 defined species. The uncharacterised proteins were incorporated into phylogenetic trees containing all ICTV RdRps from the same viral order as their best matching ICTV RdRp, by aligning sequences using MAFFT and then building trees using FastTree, both with the default settings. Monophyletic groups containing only uncharacterised proteins and viruses from a single ICTV classified order or family, with branch support ≥ 0.8 were used to make putative family and order level assignments.

### CDD Annotations

CDD annotations for uncharacterised NCBI proteins were retrieved on 2024-09-16 and assigned as: RdRp - domain only associated with RNA viral RdRp, RT-like - domain associated with reverse transcriptase or a related annotation, RNA viral non-RdRp - domain associated only with RNA viral sequences but not with RdRp, RT-like - associated with reverse transcriptase or a related protein and other - not specifically RNA viral. Assignment of the annotations was via examining the taxonomy of the sequences in which the CDD domain has previously been identified and then manually checking which annotations fell into which group and comparing these to the online descriptions of the domains. Assignments are provided in the Supplementary Data.

### Near Misses

Near misses were defined as sequences which had a HMMER hit against a viral RdRp with a score score ≥10 and ≤25. There were 1,349,599 such sequences. These sequences were compared to the nr database using DIAMOND BLASTP and filtered to keep sequences with an alignment length of >100 amino acids and an identity >40% against any sequence not labelled as uncharacterised or similar (excluding self hits). This left 577,704 sequences. A negative control dataset of similar size was generated for comparative purposes. To select the negative control sequences, a random sample of 1,000,000 unclassified sequences from the NCBI protein database was taken using the Unix shuf command, and these sequences were subjected to BLAST against nr with the same conditions as the near miss sequences, leaving 499,240 sequences. The BLAST target with the highest bit score against each query in both sets was selected.

To classify the BLAST target names into clusters, the full protein names were split into terms of two or more words, using a single space character as a delimiter. Where multiple overlapping terms were present in the same set of proteins, only the longest term was included. Terms appearing in 20 or more proteins in the near miss set were then clustered by overlap, and clusters were checked and edited manually to make biological sense, with synonyms and closely related terms grouped. Ambiguous terms were excluded. Clusters of terms and excluded terms are listed in the Supplementary Data. The number of proteins with names including any of the terms in each cluster was then counted, in the near miss and the negative control datasets. Clusters were considered to be significantly enriched if the p-value of a Fisher Exact Test (as implemented in scipy v1.10.1 [89], multiplied by the number of clusters (62) to correct for multiple testing, was less than 0.05. To calculate fold changes, a small constant of 10E-05 was added to the percentage of proteins annotated with the term, to account for zero counts in the denominator.

Details of alignments used for the near miss sequence logos are provided in the Supplementary Materials and Methods.

## Data Availability

Supplementary data has been deposited via Zenodo at https://doi.org/10.5281/zenodo.13838381. All raw data is from public repositories.

## Supporting information

Supplementary Table 1 - column labels

