## Supplementary Materials and Methods for "Uncovering hundreds of exogenous and endogenous RNA viral RdRp sequences amongst uncharacterised sequences in public protein databases"

### Filtering Reverse Transcriptase

To remove hits against reverse transcriptase from the Pfam profiles PF05919 and PF00680, which contain both RdRp and RT , a phylogenetic tree was generated consisting of the sequences included in these profiles. The alignments used were extracted from the HHsuite-3 pfamA 35.0 database [1], accessed 2023-02-17, 2021-11-23 release), and trees were generated with FastTree (v2.1.11) [2], with the default settings. New profiles were created based on the phylogeny using HMMbuild from HMMER (v3.3) [3], using subsections of the original alignment. PF05919 was divided into two subsections - RT and RdRp, while PF00680 was divided into RdRp, RT1 and RT2, as the RT section of the phylogeny had two clear subgroups of RT (Supplementary Figure 1). Ambiguously placed sequences and long branches (length >3) were excluded from the profiles. Aligned FASTA files and HMMs for the newly created profiles are available in the Supplementary Data. Sequences which originally matched either PF05919 and PF00680 were compared to each of these profiles with HMMER as for the original profiles, and assigned as either RT or RdRp based on the higher HMMER score.

### Mitovirus Phylogeny

As the mitoviruses are short and degraded, sequences were filtered to identify a shared common region of RdRp. A reliable alignment could not be generated based on only the newly identified sequences, so based on the BLAST analysis described in the main Materials and Methods section, the most similar known RdRp to each putative mitovirus RdRp was identified and these known proteins were aligned using the MAFFT linsi algorithm (v7.520) [4]. The position of the uncharacterised protein BLAST hit on its target sequence was then used to establish the position of the uncharacterised protein on the mitoviral RdRp. The 30 amino acid region spanned by the greatest number of uncharacterised proteins was identified and the uncharacterised proteins fully overlapping this region selected for phylogenetic analysis. An alignment was created incorporating these uncharacterised proteins, mitovirus reference sequences and mitoviruses identified in other virus discovery projects, sequences were aligned using the default MAFFT algorithm. The overall mitovirus phylogeny was built using FastTree2, with the default settings. Sequences were assigned to clades based on this phylogeny. Subtrees were extracted from the newick file of this phylogeny, the ginger virus phylogeny was also expanded as discussed below. The full mitovirus phylogeny shown in Supplementary Figure 4 is provided in newick format in the Supplementary Data.

To generate sequence logos, the alignment used for the overall mitovirus tree was cleaned with CIAlign (v1.1.4) [5] to remove insertions up to 1000 amino acids in length, remove terminal regions with coverage <10% or similarity <10% and remove sequences <50 amino acids in length. Sections of this alignment were then isolated based on the clade identified in the phylogeny and these sections were used directly to create the logos. Logos were drawn using CIAlign.

Tanglegrams were calculated using the Dendroscope (v3.8.10) [6] and visualised using plot_phylo (v0.0.4, github.com/KatyBrown/plot_phylo). Online HHpred [7,8] searches were performed using the server at https://toolkit.tuebingen.mpg.de/tools/hhpred against Pfam-A_v37 between 2024-04-01 and 2024-05-31.

### Ginger Mitovirus Phylogeny

A BLASTN search for the nucleotide sequence corresponding to accession KAG6467065.1 (JACMSC010000092.1 positions 112386 to 113349) against whole genome shotgun contigs from members of the Zingiberaceae family (performed using the BLAST online server on 2024-04-04) was used to identify contigs containing RdRP in other members of this family. The best scoring contig was selected for each species. ORFs in these contigs were identified using orfipy_core v0.0.4 [9] with a minimum length of 300 nt, defining ORFs as between two stop codons (between_stops=True) and allowing for partial 3' and 5' ORFs. A local BLASTP (v2.5.0) [10] search was then used to identify the best scoring ORF against KAG6467065.1 from each contig and these ORFs were aligned using MAFFT with the linsi algorithm. This alignment was cropped with CIAlign to keep only the region present in 80% of ORFs, any ORFs then less than 100 amino acids in length were discarded. The tree was created using FastTree with the default settings. The phylogeny is provided in the Supplementary Data.

### Orbivirus-Like Sequences

Initial TBLASTX searches were performed using standalone BLAST with the protein sequences identified amongst uncharacterised proteins as queries and nematode reference genomes (listed in Supplementary Data) as targets. Sequences were aligned using the MAFFT linsi algorithm and sequence similarity matrices generated with CIAlign. Chromosome-to-chromosome alignments were generated using NUCMER (v3.1) [11] with a maximum gap size of 500 and a minimum cluster size of 100. Corresponding regions were isolated using BedTools (v2.31.1) [12]. Visualisations of sequence alignments were generated using CIAlign.

RNA-seq datasets from nematodes containing potential reoviral transcripts were identified by querying the online Serratus database (accessed 2024-07-23, https://serratus.io/), [13]. All datasets were downloaded from SRA (https://www.ncbi.nlm.nih.gov/sra) between 2024-08-07 and 2024-08-14. Reads were trimmed with trim_galore (v0.6.4_dev, https://github.com/FelixKrueger/TrimGalore) then mapped sequentially to various databases to remove non-viral reads as follows: invertebrate ribosomal RNA from the SILVA database (downloaded 2020-01-15), the appropriate host genome, the human reference genome (GRCh38.p7 primary assembly) and bacterial RefSeq genomes from NCBI (downloaded 2017-10-02). Mapping was performed with bowtie2 (v2.4.1) [14], or hisat2 (for host and human) (v2.2.1) [15], both with the setting --ignore-quals and allowing 5% of the read length as mismatches. The unmapped reads after these filtering steps were assembled into contigs using the SPAdes genome assembler in rnaSPAdes mode (v3.15.5) [16]. ORFs were identified in the assembled contigs using the EMBOSS getorf function (v6.5.7.0) [17].

A set of related orbi-like proteins to use to identify further nematode orbi-like viruses was identified initially by using BLASTP against nr on the online server (accessed 2024-07-25) for the initially identified *Brugia* uncharacterised proteins. The taxonomy IDs for these sequences were identified and, for each taxon, all NCBI sequences were downloaded. All NCBI RefSeq sequences for ICTV classified members of the Orbivirus genus were also included. The Pfam profiles for Orbivirus proteins (listed in the Supplementary Data) were used as targets for HHsearch (v3.3.0) [1] with the default settings, to classify which viral proteins these sequences correspond to. The proteins, their host species and their matching orbivirus profiles are listed in the Supplementary Data.

To identify RdRp in the assembled SRA contigs, they were compared to VP1 proteins from this set using DIAMOND BLASTP (v0.9.14) [18], with the default settings. For *H. contortus*, which has intact ORFs, the highest scoring RdRps from these results were used for all further analysis. For other nematodes, the identified contigs were used as input to miniprot below to reconstruct ORFs.

Miniprot (v0.13-r248) [19] was used to reconstruct putative ORFs from degraded EVE sequences with relaxed settings as follows (as the sequences are expected to differ substantially from the reference) - no splicing (-S), kmer size (-k) and kmer size for second round of chaining (-l) 6, modimisers bit (seed density, -M) 0, frameshift penalty (-F) 6, minimum number of syncmers (-n) 6, minimum proportion of query aligned (--outc) 0.1. Chromosome regions identified above and the non-*H. contortus* SRA contigs were used as queries and the known orbivirus VP1 proteins described above were initially used as target sequences. The process was repeated iteratively three times, with the ORFs resulting from the previous round as queries, to identify additional or longer regions. Only regions with at least 100 matching nucleotides were retained. Reconstructed ORFs are available in the Supplementary Data, as is the information required to regenerate these ORFs from the contigs. For SRA contigs, the contig sequences which contained the ORFs are also available in the Supplementary Data.

To identify additional segments in *H. contortus*, HMMscan and HHpred were used as described above for all *H. contortus* ORFs against the Orbivirus Pfam profiles, plus against additional profiles generated by clustering (using MMSeqs with at 35% similarity and 20% coverage in cluster mode 0) the unidentified proteins from known orbi-like viruses. Proteins scoring >30 with either of these approaches were then checked with online BLAST and compared using ESMfold and FoldSeek. For ESMfold, proteins were split into substrings with a length of 400 amino acids and an overlap of 200 amino acids. These sequences were folded using the ESM Fold Sequence online server (accessed via API on 2024-09-10 at https://api.esmatlas.com/foldSequence/v1 [20]. Similarity scores were then calculated using FoldSeek (v6.29e2557), [21] against structures generated using ESMfold, with the same settings, for the known orbi-like proteins. Sequences with a FoldSeek bit score >50 were then additionally verified using the HHpred online server against Pfam-A_v37 (accessed 2024-09-13).

### Aligning Near-Miss Proteins

For the top ten annotations identified as enriched amongst near-miss proteins compared to the controls, BLAST target sequences matching near-miss proteins were compared directly to the relevant HMM profiles using HMMscan. The matching regions identified by

HMMscan were extracted from both the profile alignments and the target sequence alignments. New HMMs were generated using just these regions of the profile alignments using HMMbuild and the target sequence regions were realigned to these HMMs using HMMalign with the --mapali setting. Sequence logos were generated using CIAlign.
