## Supplementary Figure Legends for "Uncovering hundreds of exogenous and endogenous RNA viral RdRp sequences amongst uncharacterised sequences in public protein databases"

**Supplementary Figure Legends**
**Supplementary Figure 1**
Approximate maximum-likelihood [1] phylogenetic trees for the sequences from Pfam profiles PF00680 (A) and PF05919 (B). Sequences labelled in their original database as RdRp are shown in blue and those labelled as RT or similar are shown in orange, unclassified proteins are shown in green, proteins classified as non viral are shown in purple. Specific mislabelled proteins mentioned in the text are highlighted. Clades containing only named RdRp or RT and uncharacterised proteins have been collapsed in some cases for clarity; full trees are available in the Supplementary Data.

**Supplementary Figure 2**
Left: Plot showing the regions of RdRp represented in mitovirus-like EVEs from different plant orders, based on BLAST [2] comparison to known mitoviral RdRps. Vertical lines represent the 30 amino acid region which was most commonly included. Right: As for the left plot but including only sequences which did fully overlap the 30 amino acid common region.

**Supplementary Figure 3**
Left: Plot showing the regions of RdRp represented in mitovirus-like EVEs from non-plant taxa, based on BLAST [2] comparison to known mitoviral RdRps. Vertical lines represent the 30 amino acid region which was most commonly included. Right: As for the left plot but including only sequences which did fully overlap the 30 amino acid common region.

**Supplementary Figure 4**
Approximate maximum-likelihood [1] phylogenetic tree for known members of the Mitoviridae and newly identified related sequences. Tip labels and tip label text colours provide the phylogenetic group to which the sequence was originally labelled - non-plant sequences are assigned as fungi (purple), metagenomic (dark blue), viruses (orange), metazoa (pink), bacteria (yellow), SAR (cyan), Discoba (light blue) or unclassified (grey). Plant sequences are shown in green and labelled with the plant order in which they were identified. Previously known sequences are labelled with their GenBank accession where available and otherwise another unique identifier from their source database or study. Sequences are labelled with their source database or study, these are provided as a list in the Supplementary Data. Newly identified sequences are in italics if they have a DNA source, otherwise they have an RNA source. Clades are labelled with their apparent mitoviral genus, the Duamitovirus genus is subdivided into the P, F1 and F2 clades.

**Supplementary Figure 5**
Sequence logos for RdRp motifs A to C from the Duamitovirus P, F1 and F2 clades and from the genera Unuamitovirus and Triamitovirus and Kvaramitovirus. Variable positions are labelled in red and with “$”. Conserved positions are labelled in blue

and with “*”. Positions are relative to the full length alignment available in the Supplementary Data.

**Supplementary Figure 6**
Specific subclades of the OTU phylogenetic trees showing EVEs of interest. (A) Kitaviridae and Virgaviridae like arthropod EVEs, (B) cytorhabdo-like EVEs from *Pomphorhynchus laevis*, (C) lepidopteran rhabo-like EVEs, (D) *Perkinsus olseni* arli-like sequences, (E) lepidopteran orthomyxo-like EVEs, (F) *Coccinella septempunctata* durna-like EVEs, (G) *Dinoponera quadriceps* partiti-like EVEs, (H) arthropod and nematode orbi-like EVEs. Newly identified sequences are coloured and labelled, DNA-source sequences are blue and labelled with “*”, RNA source sequences are purple and labelled with “**†”.** Previously known sequences are labelled with their GenBank accession where available and otherwise another unique identifier from their source database or study. Sequences are labelled with their source database or study, these are provided as a list in the Supplementary Data.

**Supplementary Figure 7**
CIAlign [3] mini alignment showing the alignment between fragmented regions of a *Perkinsus olseni* arli-like EVE to recreate the putative full length regions. The alignment is available in the Supplementary Data.

**Supplementary Figure 8**
CIAlign [3] mini alignment showing the alignment between the six transcript isoforms at locus LOC123308761 in *Coccinella septempunctata*. ORFs from annotated proteins from these transcripts are shown with red lines if they contained the partiti-like EVE and blue lines if they did not. The green vertical line indicates two regions which show some homology with each other and may have caused misannotation.

**Supplementary Figure 9**
(A, C) Approximate maximum-likelihood [1] phylogenetic trees for the segments encoding VP3 (A) and VP4 (C) in the orbi-like likely exogenous virus identified in *Haemonconchus contortus*. Likely complete sequences have the suffix “c” and likely partial sequences have the suffix “p”. Newly identified sequences are shown in pink and labelled by their SRA accession, known sequences are labelled with their GenBank accession and organism name. (B, D)The distribution of ORF lengths of previously annotated VP3 (B) or VP4 (D) proteins in known members of the orbi-like clade (black dots) compared to those identified in the *H. contortus* contigs (pink line, full-length sequences only).

**Supplementary Table Legends**
**Supplementary Table 1**
Table containing full details of the 3,560 RdRp-like sequences identified in this study, including metadata for the original NCBI / UniProt protein records, assigned virus

taxonomic groups, results from screening with HMMER, BLAST, FoldSeek, PalmScan and results from the CDD and known virus analyses. Column labels are listed in the accompanying file ST1_column_labels.tsv.


1. Price, M. N., Dehal, P. S., & Arkin, A. P. (2010). FastTree 2--approximately maximum-likelihood trees for large alignments. *PloS one*, *5*(3), e9490. https://doi.org/10.1371/journal.pone.0009490

2. Altschul, S. F., Gish, W., Miller, W., Myers, E. W., & Lipman, D. J. (1990). Basic local alignment search tool. *Journal of molecular biology*, *215*(3), 403–410. https://doi.org/10.1016/S0022-2836(05)80360-2

3. Tumescheit, C., Firth, A. E., & Brown, K. (2022). CIAlign: A highly customisable command line tool to clean, interpret and visualise multiple sequence alignments. *PeerJ*, *10*, e12983. https://doi.org/10.7717/peerj.12983
