## Supplementary figures and images for "Uncovering hundreds of exogenous and endogenous RNA viral RdRp sequences amongst uncharacterised sequences in public protein databases"

### Supplementary Figure 1

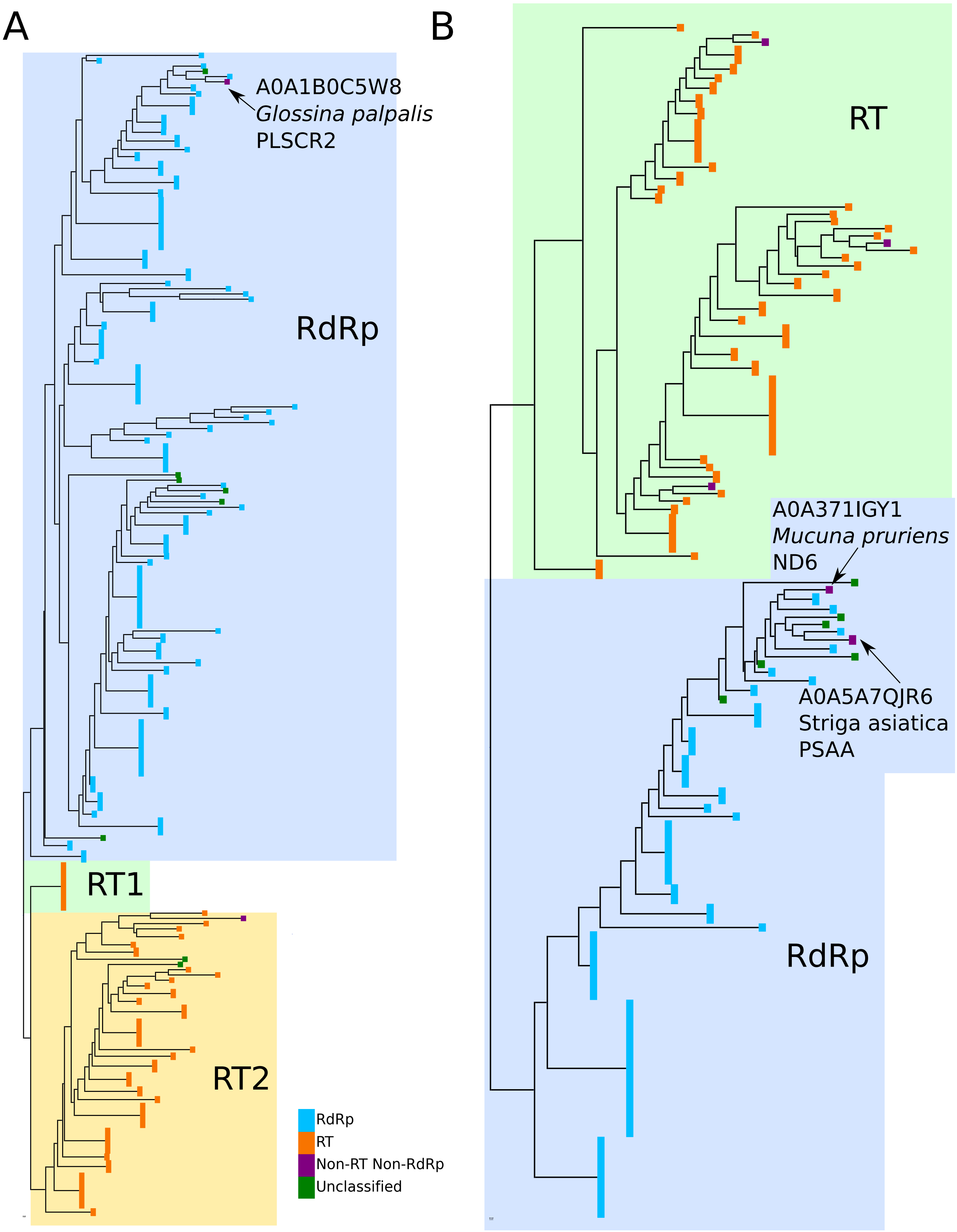

### Supplementary Figure 2

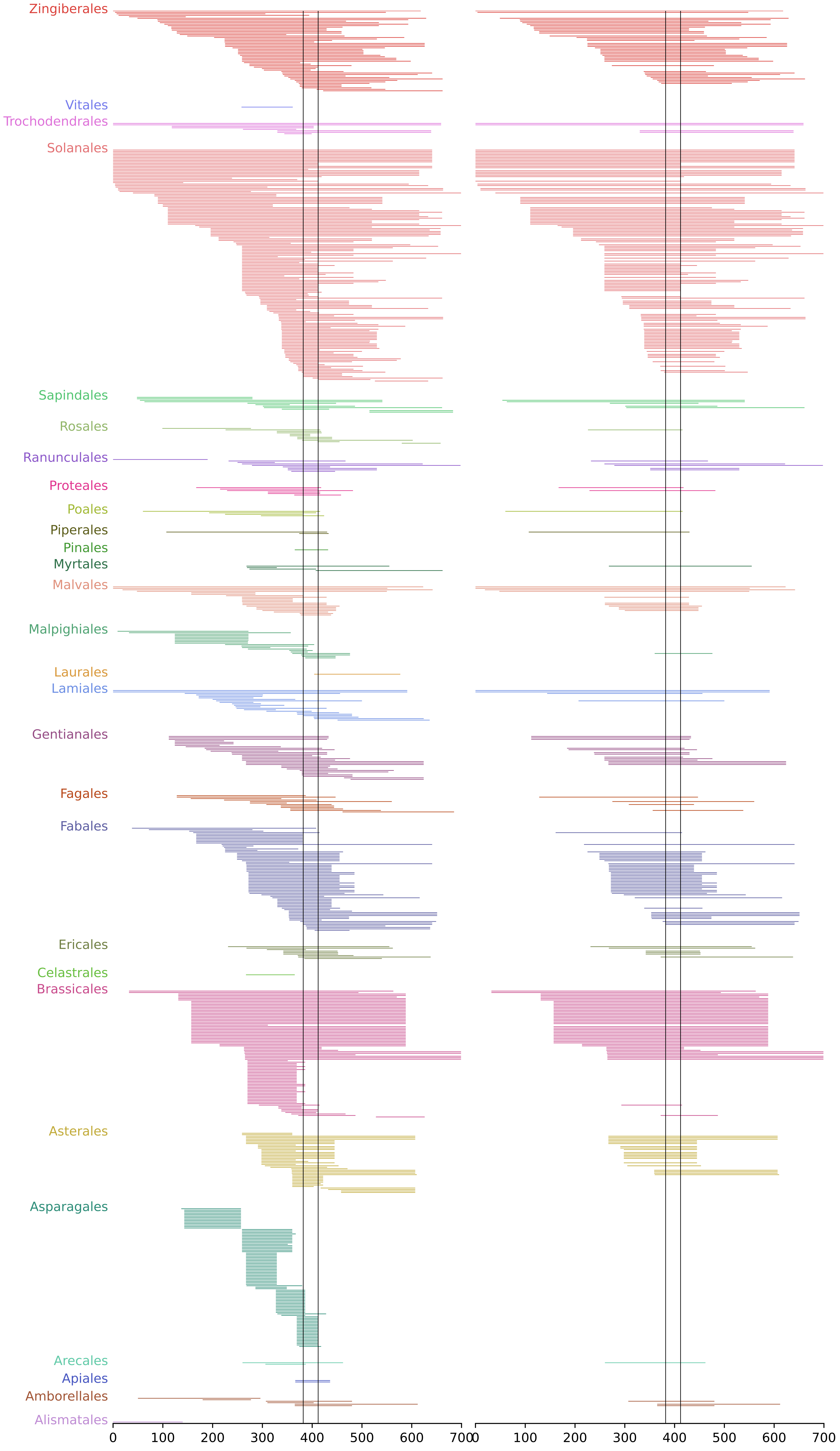

### Supplementary Figure 3

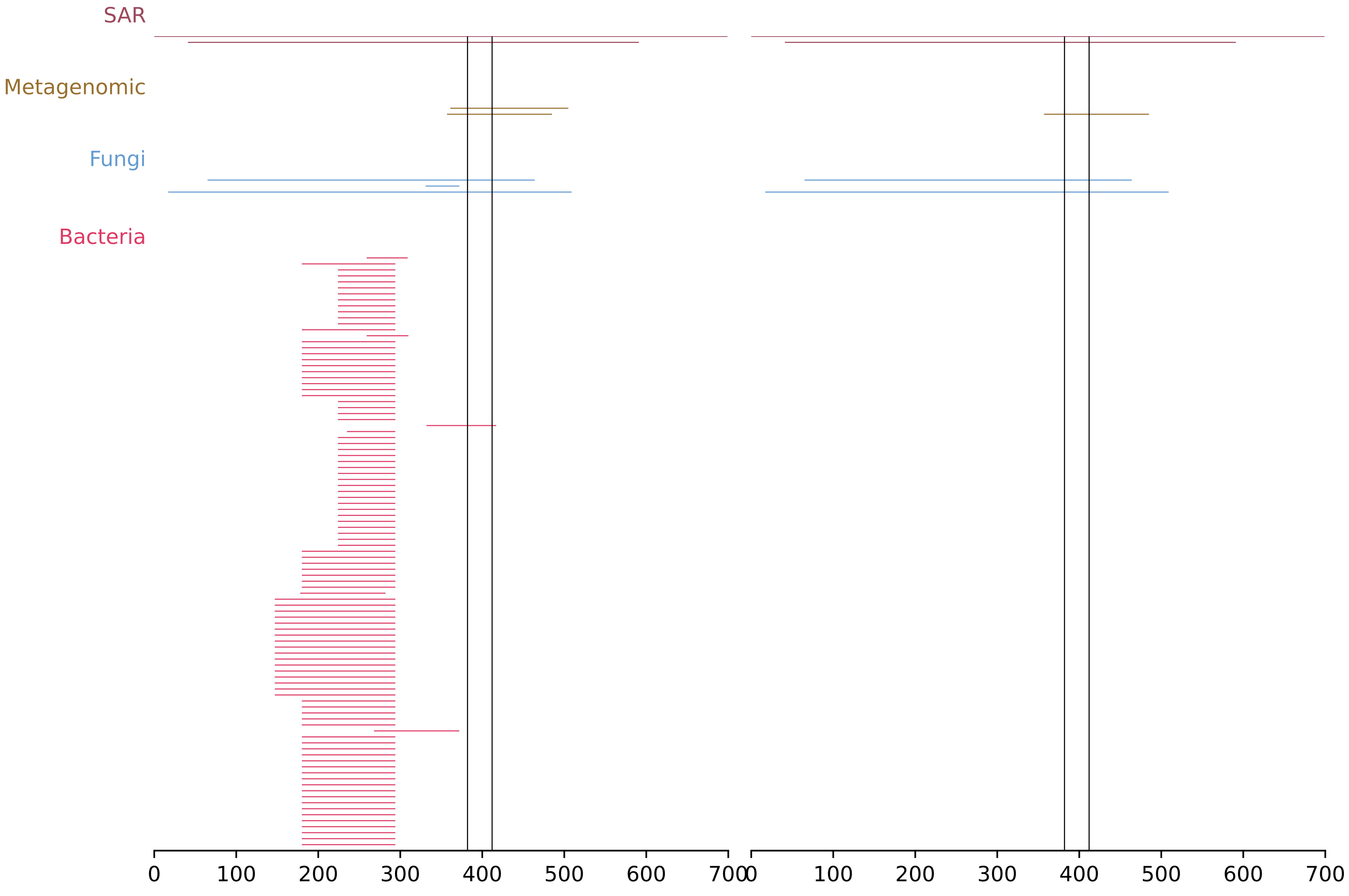

### Supplementary Figure 5

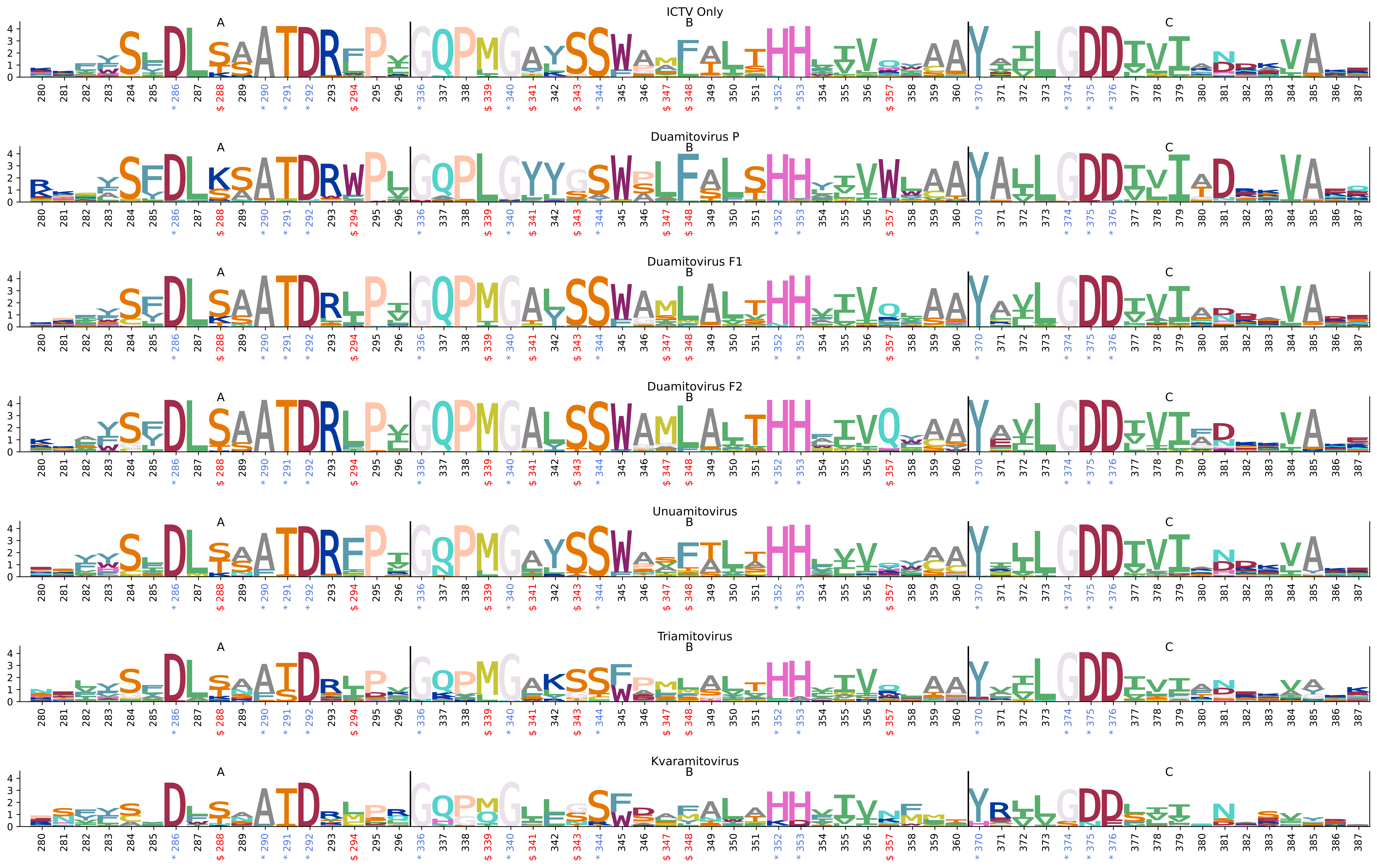

### Supplementary Figure 6

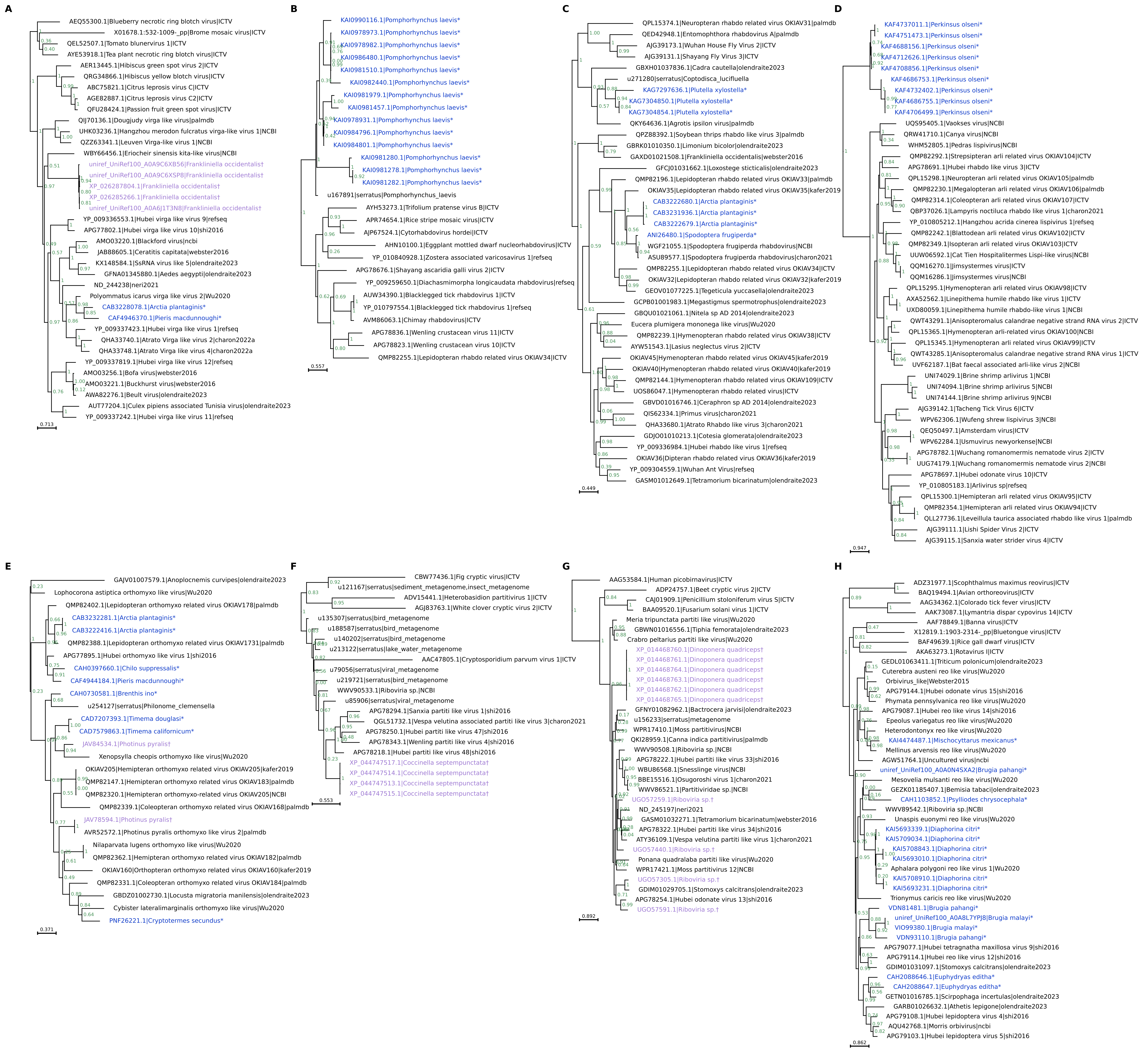

### Supplementary Figure 7

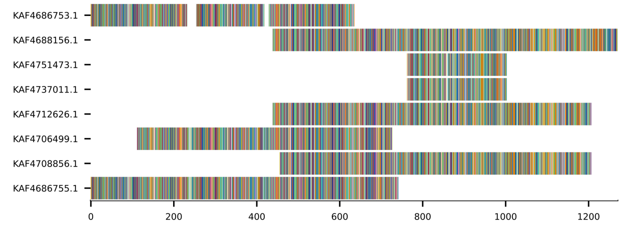

### Supplementary Figure 8

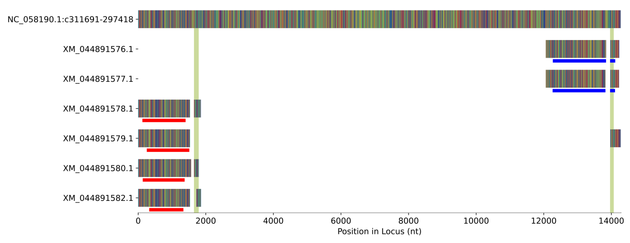

### Supplementary Figure 9

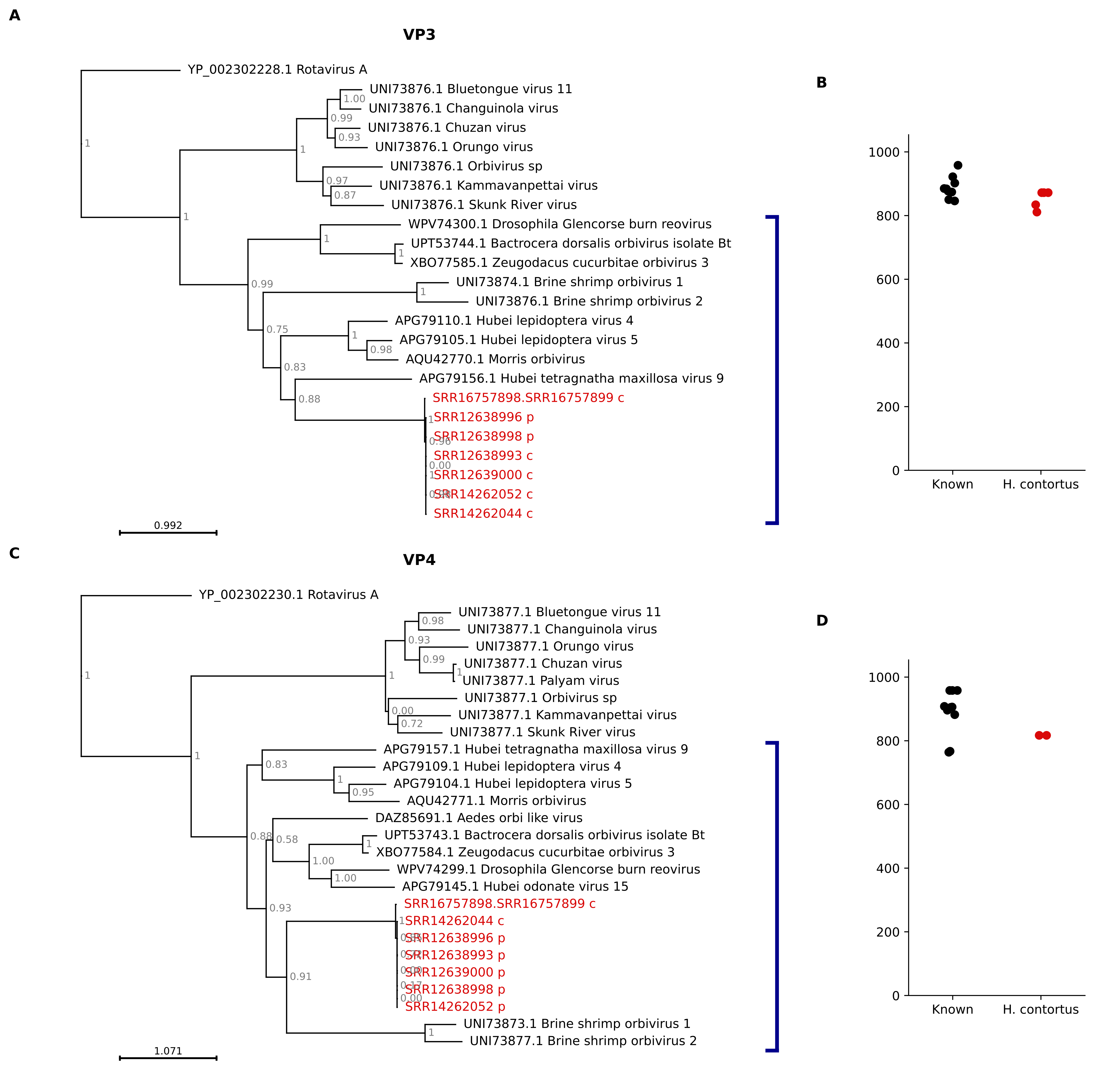
